## Supplementary Figures 1 to 7, Supplementary Tables 1 to 14, Supplementary Note 1 and Supplementary References for "ERK1/2 is an ancestral organising signal in spiral cleavage"

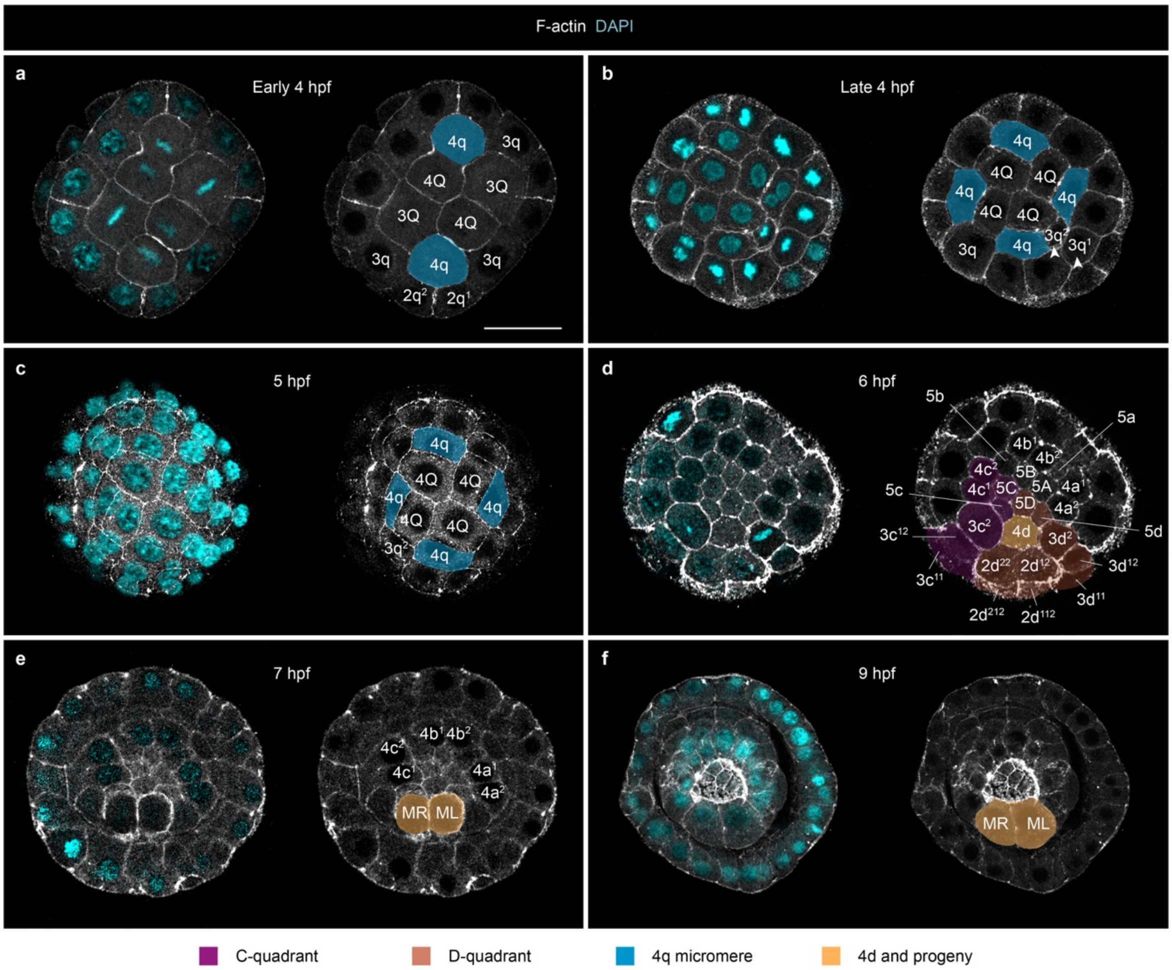

**Supplementary Figure 1. Dynamics of cell division of the fourth micromere quartet in *O. fusiformis*. a–f** z-stack confocal projections of the vegetal pole of whole mount embryos stained with phalloidin (F-actin, gray; it shows cell boundaries) and DAPI (nuclei) during the onset and early division of the fourth micromere quartets and the blastomere inferred to be the 4d cell based on ERK1/2 activity. The fourth micromere quartet emerges asymmetrically (early 4 hours post fertilisation, hpf), which is soon followed by the asymmetric division of one of the 3q micromeres (arrowheads) (late 4 hpf). The vegetal pole is however quatri-radial at 5 hpf, with the 4d arresting its division with respect to 4a–c, and only cleaving into MR and ML (4d daughter cells) after ingression at 7 hpf. False colouring indicates cell or quadrant identities. Scale bar is 50 µm.

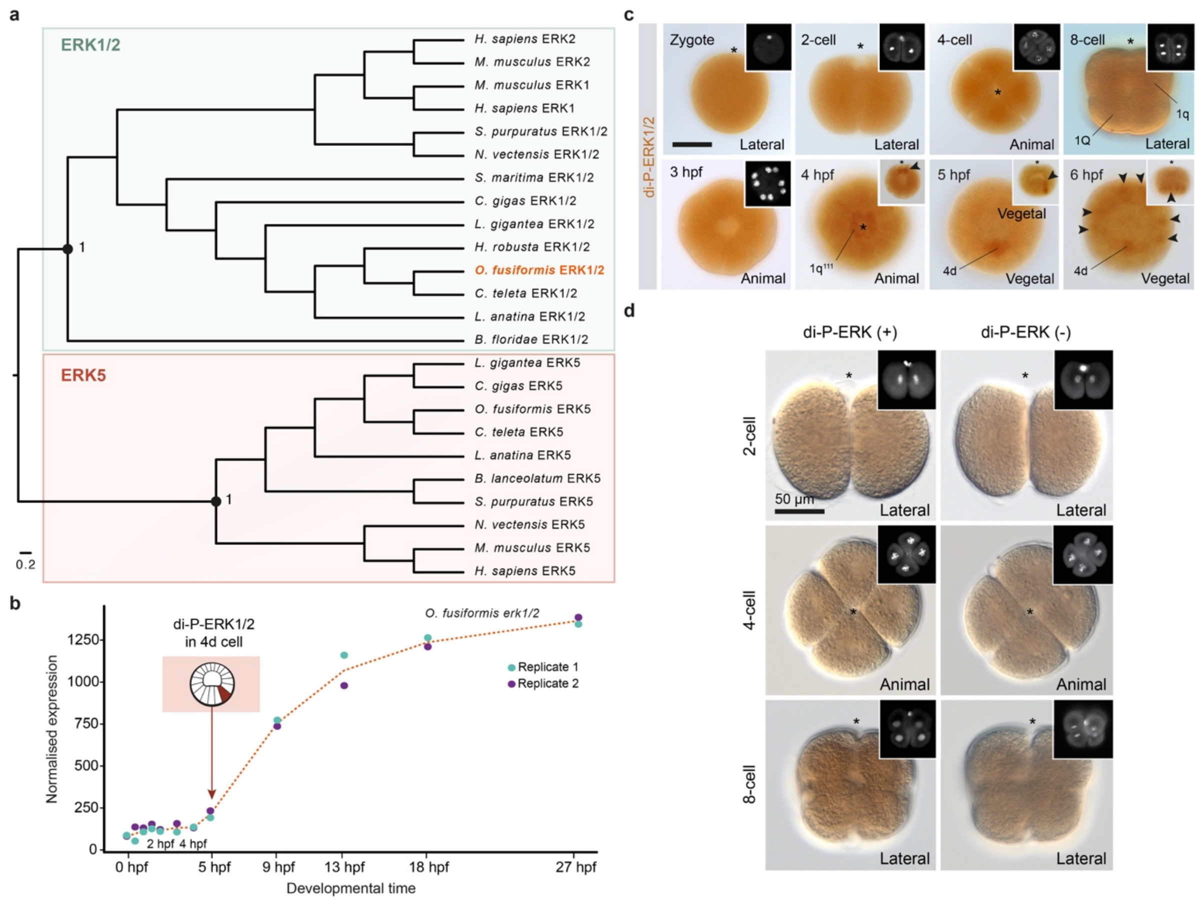

**Supplementary Figure 2. ERK1/2 activity during *O. fusiformis* early embryogenesis. a** Maximum likelihood orthology assignment for *O. fusiformis* ERK1/2*,* using ERK5 as outgroup. Only bootstrap values for major nodes are shown. **b** Normalised expression of *erk1/2* during *O. fusiformis* development, from the active oocyte to the mitraria larval stage in hours post fertilisation (hpf). The time of the specification of 4d is highlighted with a schematic drawing. Coloured dots indicate values of expression for each replicate. The dotted red line shows the mean value of expression. *erk1/2* expression starts to increase at 4 hpf, which is consistent with the enrichment of di-phosphorylated-ERK1/2 in specific blastomeres from 4 hpf onwards (see Figure 2). **c** Whole mount immunohistochemistry against di-phosphorylated-ERK1/2 (di-P-ERK1/2; dark orange) during spiral cleavage (from the zygote to 6 hpf) in *O. fusiformis*. Low background levels are detected during early divisions (zygote to 3 hpf). Later, di-P-ERK1/2 is enriched in the four $1q^{111}$ animal micromeres at 4 hpf, the 4d micromere at 5 hpf and in six vegetal cells forming a bilaterally symmetrical pattern (arrowheads) plus 4d at 6 hpf. Insets show either nuclear staining (gray; zygote to 3 hpf) or lateral views (4 to 6 hpf). **d** Whole mount immunohistochemistry against di-P-ERK1/2 and the corresponding negative control (without primary antibody). Background levels are comparable between the two conditions. Insets show nuclear staining (gray). In **c** and **d**, asterisks point to the animal pole.

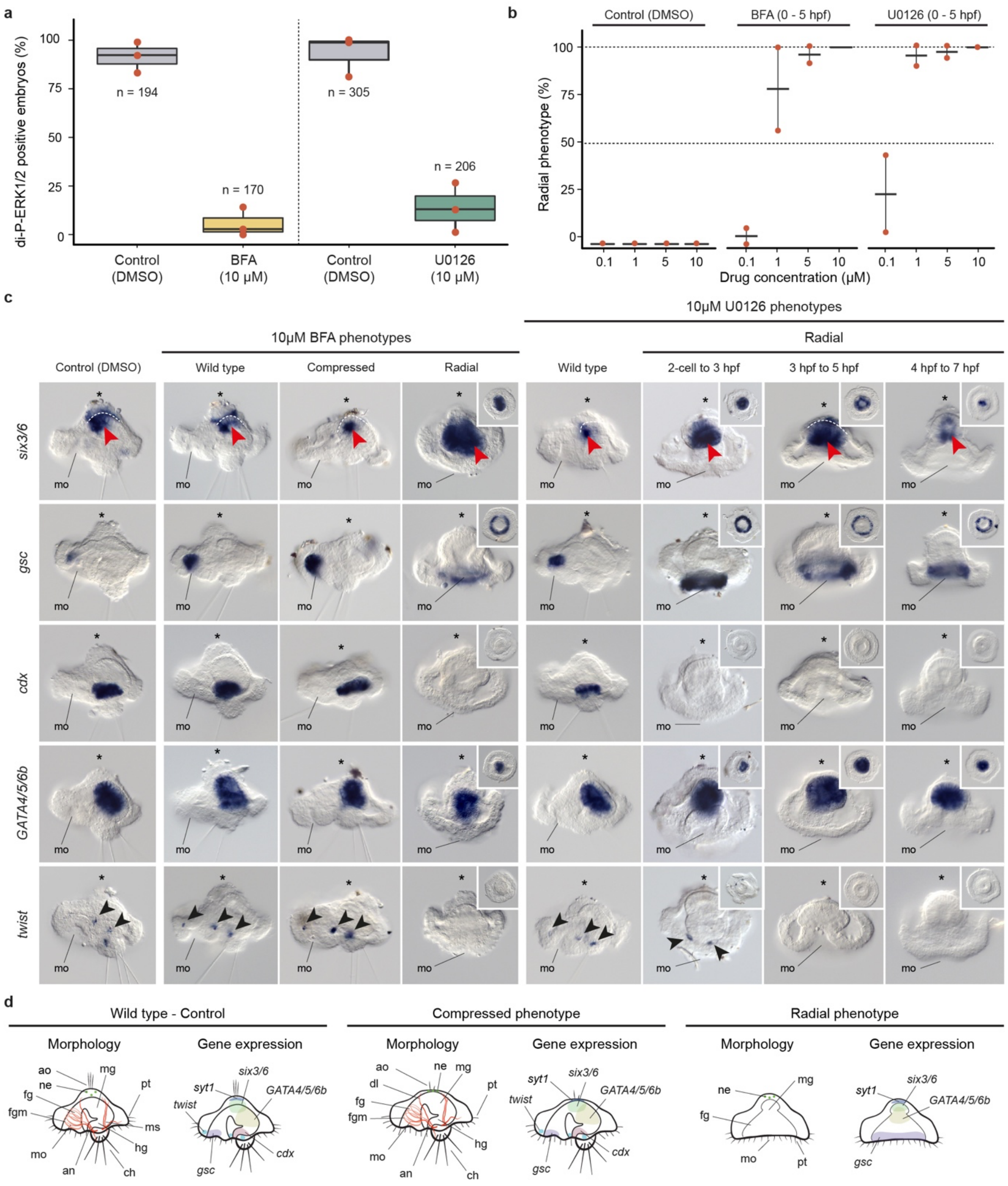

**Supplementary Figure 3. The effect of BFA and U0126 treatment in *O. fusiformis* development. a** Boxplots depicting the percentage of coeloblastulae showing di-phosphorylated ERK1/2 (di-P-ERK1/2) enrichment in the 4d micromere after 0.5 to 5 hours post fertilisation (hpf) treatment with brefeldin A (BFA), U0126 or DMSO (control). Red dots indicate the values for each experimental replicate. **b** Plot indicating the percentage of radial phenotypes obtained when treating embryos from 0.5 to 5 hpf with a range of BFA and

U0126 concentrations. A concentration of 10 µM shows the greatest penetrance and was thus used throughout the study. **c** Whole mount *in situ* hybridisation of apical (*six3/6*), oral (*gsc*), posterior/hindgut (*cdx*), midgut (*GATA4/5/6b*) and trunk mesodermal (*twist*) gene markers in control and treated embryos fixed at larval stage. Radial phenotypes have a reduction of the apical marker (although *six3/6* endodermal expression is maintained and/or expanded; red arrowheads; the dotted white line separates ectodermal from endodermal expression), radial expansion of an oral gene, loss of posterior structures (chaetae) and posterior/trunk mesoderm gene expression and retain the expression of a midgut gene. Radial larvae treated with U0126 from the 2-cell stage to 3 hpf show scattered *twist* expression in some embryos (black arrowheads) and embryos treated from 4 to 6 hpf are seemingly more elongated along the apical-oral axis. The compressed larval phenotype after BFA treatment from 0.5 to 4 hpf have a reduced apical organ and apical tuft (first row) and reduced expression of apical markers (*six3/6*), but otherwise normal morphology besides an obliterated internal blastocele. Asterisks indicate the apical pole. Insets are ventral views. **d** Schematic drawings (not to scale) of the control and treated larval phenotypes, depicting the morphological landmarks, genes and gene expression patterns considered to ascribe treatment outcomes to each of the phenotypic categories (summarised in Supp. Table 3). an: anus; ao: apical organ; ch: chaetae; dl: dorsal levator muscles; fg: foregut; fgm: foregut muscles; hg: hindgut; mg: midgut; mo: mouth; ms: muscles; ne: neurons; pt: prototroch.

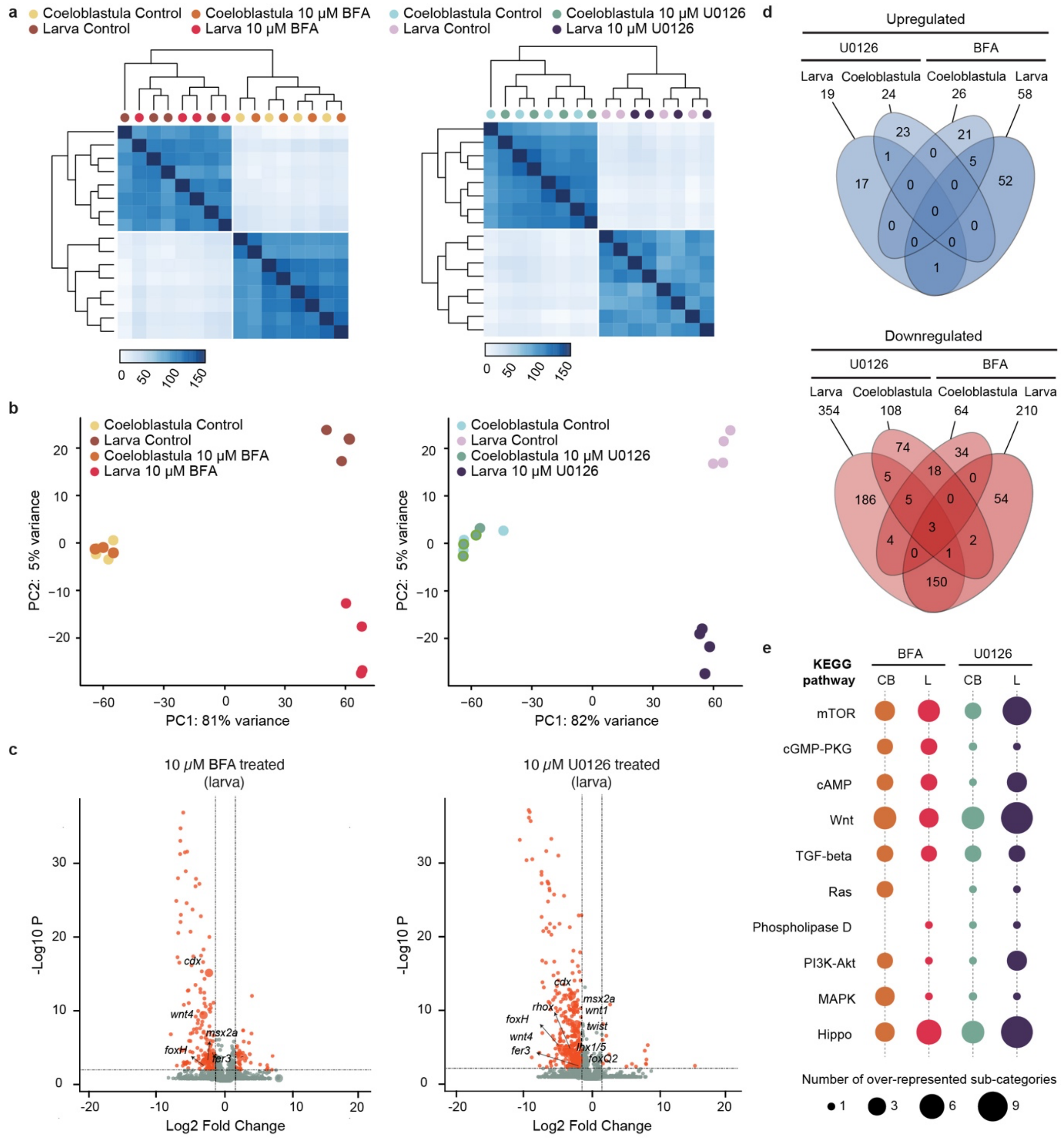

**Supplementary Figure 4. Differential expression analyses in BFA and U0126 treated embryos. a** Hierarchically clustered pairwise correlation matrix between brefeldin A (BFA) and U0126 treated and control samples. All replicates of the same condition are highly correlated. **b** Principal component (PC) analysis plot for BFA (left) and U0126 (right) RNA-seq analyses. Coeloblastula and larval samples are clearly separated, as well as treated and control samples fixed at larval stage, which reflects the increase in the number of differentially expressed genes at this later time point compared to the coeloblastula stages (as shown in panel **d**). **c** Volcano plots for BFA and U0126 treated embryos studied at 24hpf (larval stage). Red dots show differentially expressed (DE) genes and analysed candidate genes are labelled in each comparison. In both conditions, most DE genes are downregulated. **d** Venn diagrams showing the number of up- (top) and downregulated (bottom) DE genes shared between different conditions. **e** Ten most overrepresented KEGG pathways in each of the four conditions analysed by RNA-seq. The size of the coloured dots is proportional to the number of overrepresented sub-categories nested into a given KEGG pathway.

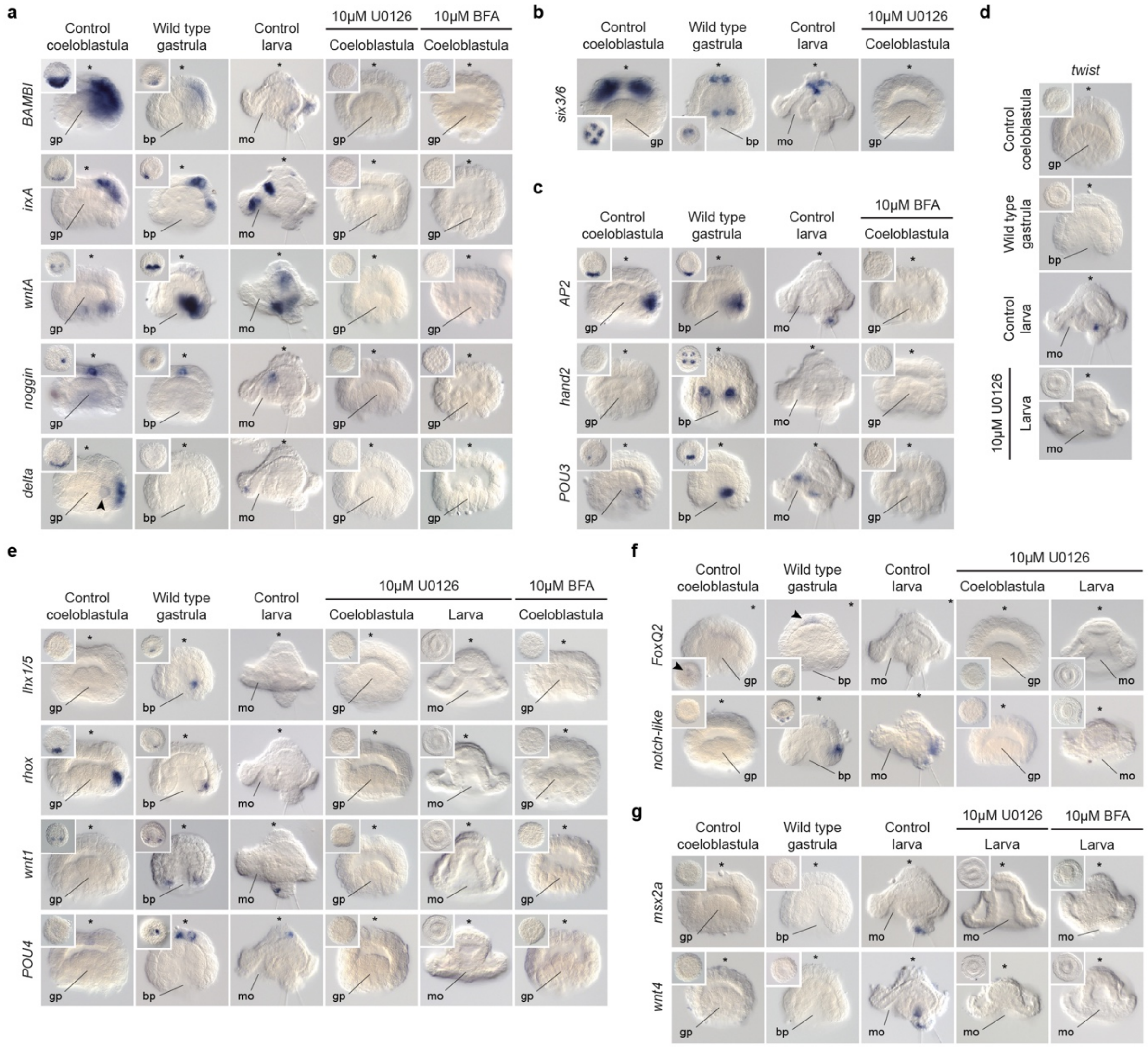

**Supplementary Figure 5. Validation of the differential expression analyses. a**–**g** Whole mount *in situ* hybridisation of control and treated embryos, as well as wild type gastrula stage to reconstruct the time course of expression of each gene. **a** Candidate genes differentially expressed at the coeloblastula stage in both BFA and U0126 treatments. *BAMBI*, *irxA* and *delta* are expressed in the dorsal ectoderm and 4d (only *delta*) at the coeloblastula stage. *WntA* is expressed in the posterior blastoporal lip and probably larval mesoderm, and *noggin* is detected in the apical ectoderm of the blastula and gastrula. **b** *six3/6* (expressed at the apical ectoderm and larval oesophagus) is differentially expressed at the coeloblastula stage in U0126 treatment only. **c** Candidate genes differentially expressed at the coeloblastula stage in BFA treatment only. *AP2* is expressed in posterodorsal ectoderm, and *hand2* and *POU3*

are likely expressed in mesodermal cell types. **d** *twist* (expressed in mesoderm) is differentially expressed at the larval stage in U0126 treatment only. **e** Candidate genes differentially expressed at the coeloblastula stage in both BFA and U0126 treatments, as well as at the larval stage in U0126 treatment. *Lhx1/5* and *rhox* are expressed in mesodermal cells, *wnt1* is detected in posterior ectodermal cells and *POU4* in apical ectodermal cells. **f** Candidate genes differentially expressed at the coeloblastula and larval stages in U0126 treatment. *foxQ2* is expressed in the apical ectoderm and *notch-like* is detected in the posterodorsal ectoderm of the gastrula and larva. **g** Candidate genes differentially expressed at the larval stage in both BFA and U0126 treatments. *Msx2a* is expressed in the posterior ectoderm and *wnt4* in posterior mesoderm. In all cases, the expression patterns disappear in BFA and U0126 treated embryos compared to the control condition at the relevant stages. Main panels are lateral views and insets are ventral views (except for genes expressed apically, where insets are apical views). Asterisk point to the animal/apical pole. Arrowheads point to 4d in **a** and to the apical organ in **f**. bp: blastopore; gp: gastral plate; mo: mouth.

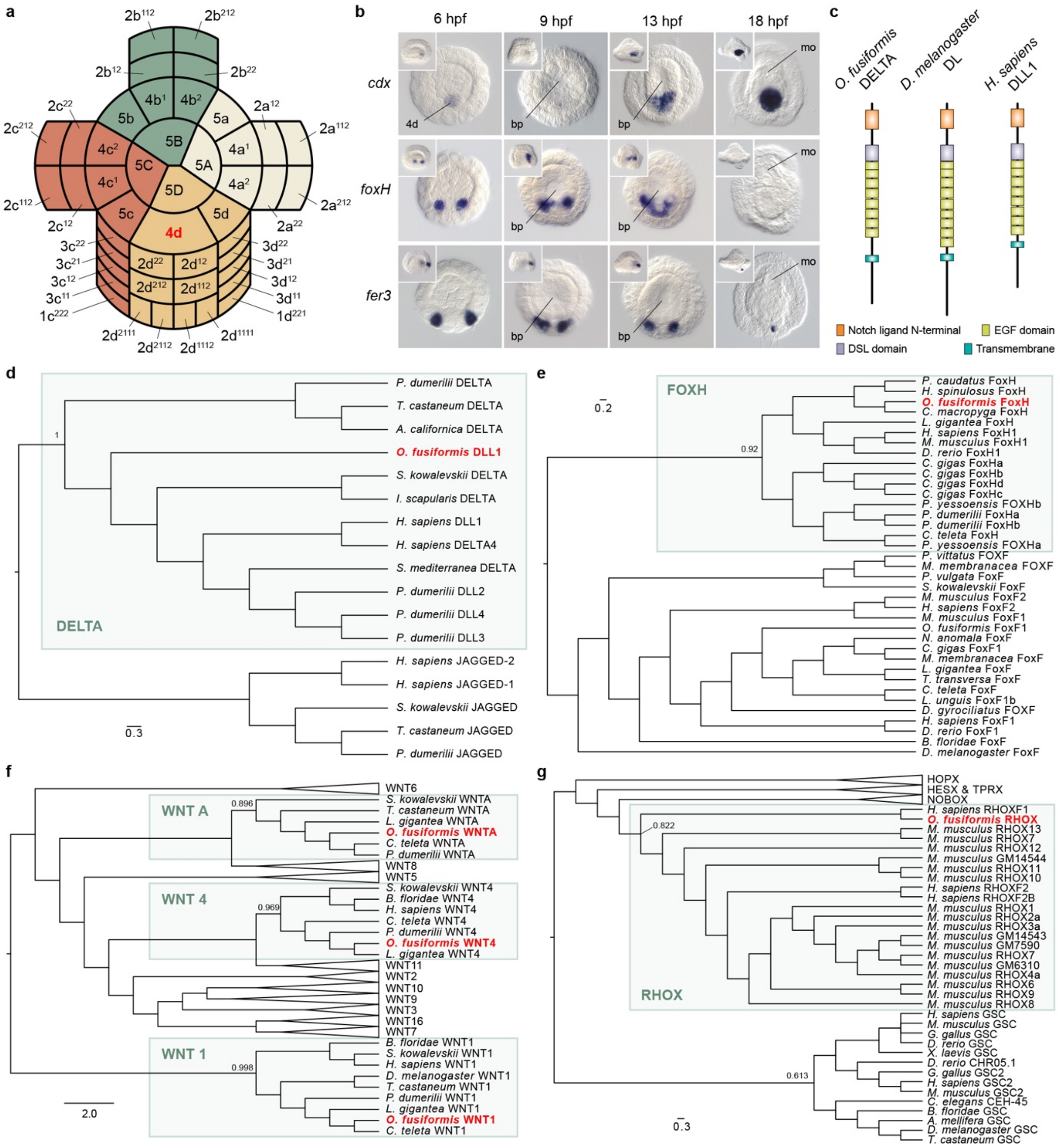

**Supplementary Figure 6. Genes patterning the D-quadrant after the specification of the 4d-organiser cell. a** Schematic drawing of the cellular arrangements and cell nomenclature of the vegetal pole just after the specification of the 4d micromere in *O. fusiformis* embryo at 5.5 hours post fertilisation (hpf). Each colour depicts a different quadrant. **b** Ventral views of the time course of expression of *cdx*, *foxH* and *fer3* via whole mount *in situ* hybridisation from 6 hpf to 18 hpf (early larval stage) during *O. fusiformis* embryogenesis (insets show lateral views). While *cdx* is expressed in the cells forming the hindgut, *foxH* appears to upregulate at the earliest stages of cells giving rise to mesodermal derivatives. The gene *fer3* is expressed in two posterior most cells of unknown function in the gastrula and larva. Insets are lateral views. **c** Schematic drawing of the protein domain architecture of the Notch ligand DELTA in *O. fusiformis* in comparisons with the domain structures of the Delta orthologs in *D. melanogaster* and *H. sapiens*. **d–g** Maximum likelihood orthology assignments of DELTA, FOXH, WNT ligands, and RHOX in *O. fusiformis*. Only bootstrap values supporting each major clade are shown. In **b**, bp stands for blastopore and mo for mouth.

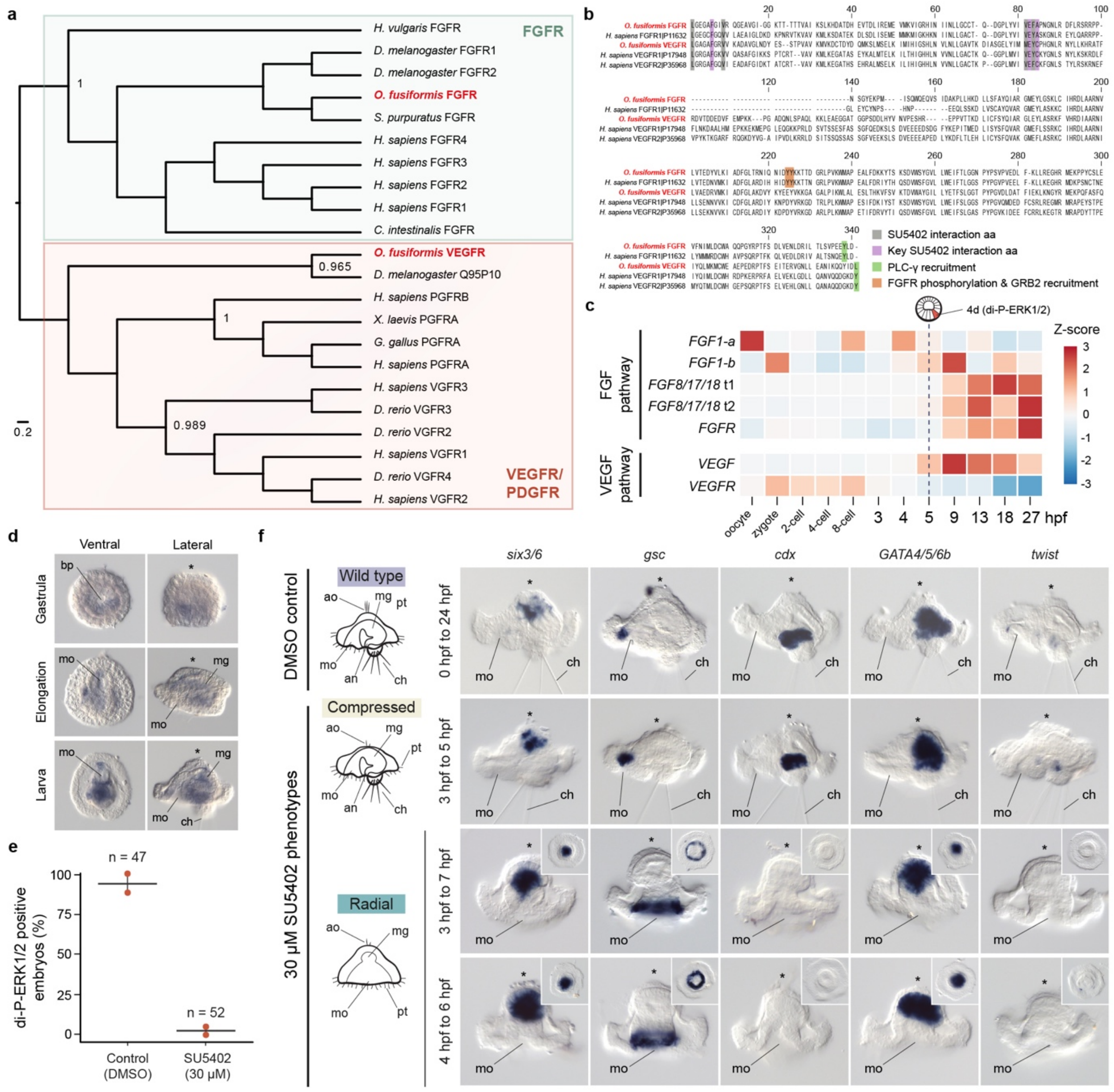

**Supplementary Figure 7. The FGF pathway and FGFR activity during 4d specification in *O. fusiformis*. a** Maximum likelihood orthology assignment of fibroblast growth factor receptor (FGFR) and vascular endothelial growth factor receptor (VEGFR). Only bootstrap values for key nodes are shown. **b** Multiple protein alignment between *O. fusiformis* and *H. sapiens* FGFR and VEGFR orthologs indicating conserved residues for the interaction of the drug SU5402 with these receptors. **c** Temporal time course of expression of members of the FGF and VEGF pathways in *O. fusiformis*. The heatmap depicts the normalised z-score value of expression for each gene. The vertical dotted line highlights the specification of the 4d micromere. **d** Whole mount *in situ* hybridisation time course of expression of *fgfr* from gastrula to larval stages in *O. fusiformis*. At the gastrula stage, *fgfr* is expressed in the posterior archenteron wall. During organogenesis and elongation, as well as at the larval stage, *fgfr* is detected in putative mesodermal tissues and midgut. **e** Plot depicting the proportion of treated and control embryos exhibiting di-phosphorylated ERK1/2 enrichment in the 4d cell at 5 hours post fertilisation (hpf) after SU5402 drug treatment from 0.5 to 5 hpf. **f** Morphological and molecular characterisation of SU5402 treated embryos fixed at the larval stage. The first column shows diagrams of lateral views of the wild type, compressed and radial phenotypes. From the second column onwards, whole mount *in situ* hybridisations of control and SU5402 treated conditions. The "compressed" larvae are similar to brefeldin A (BFA) "compressed" larvae, with reduced apical organ and apical tuft (second row) and reduced expression of apical markers (*six3/6*), but other tissues are not affected. Radial larvae are phenocopies of BFA and U0126 radial larvae with reduction of apical markers (*six3/6*) in the apical ectoderm, radial expansion of oral genes (*gsc*), loss of posterior structures (chaetae) and expression of posterior endomesodermal markers (*cdx*, *twist*), but normal expression of an endodermal gene (*GATA4/5/6b*). Insets are ventral views. Asterisks point to the animal/apical pole. ao: apical organ; an: anus; bp: blastopore; ch: chaetae; gp: gastral plate; mg: midgut; mo: mouth; pt: prototroch.

### Supplementary Table 1. The role of di-P-ERK1/2 in spiralians

| Reference | Clade | Species | Type of cleavage | Description of di-P-ERK1/2 immunoreactivity during early development | Axial defects after MAPK inhibition | U0126 concentration | Phenotype |
|---|---|---|---|---|---|---|---|
| REF[1] | Cheilostomatida (Bryozoa) | *Membranipora membranacea* | Non spiral (biradial) | 3D | Yes | 1,10 or 25 µM | But, presumably maternal, prior to the activation in 3D. |
| REF[2] | Gastropoda (Mollusca) | *Haliotis asinina* | Equal | 3D | Yes | 10 or 50 µM | Reduced dorsal and ventral tissue and no torsion at lower concentrations. Post trochal protrusion with no evident ventral, dorsal tissue o posterior. |
| REF[3] | Gastropoda (Mollusca) | *Tritia obsoleta* | Unequal | 3D + the micromeres known to require a signal from 3D | Yes | 10 µM | Post trochal protrusion with no evident ventral, dorsal tissue o posterior. Lack of eyes and velar lobes. |
| REF[4] | Gastropoda (Mollusca) | *Tectura scutum* | Equal | 3D | Yes | 10 or 50 µM | Loss or reduction of larval retractor muscles (derived from 4d) at lower concentration. No shell or operculum at higher concentrations. |
| REF[5] | Gastropoda (Mollusca) | *Crepidula fornicata* | Equal | Initially in the progeny of the first quartet micromeres, just prior to the birth of the third quartet (e.g., late during the 16-cell and subsequently during the 20-cell stages). Afterwards,in 3D just prior to the 24-cell stage, transiently in 4d and finally in a subset of animal micromeres immediately following those stages. | Yes | 10 or 25 µM | Gastrulation defects. 1st, 2nd and 3rd quartet micromeres form protrusion with no evident ventral or dorsal tissue. No eyes or shell. |
| REF[6] | Gastropoda (Mollusca) | *Patella vulgata* | Equal | 3D | No | 10 or 50 µM | Changes in the expression of brachyury, but ventral and dorsal tissues are evident |
| REF[7] | Gastropoda (Mollusca) | *Testudinalia testudinalis* | Equal | 3D | Unclear | 10 or 40 µM | |
| REF[4] | Gastropoda (Mollusca) | *Lymnaea palustris* | Equal | 3D | Not studied | | |
| REF[4] | Polyplacophora (Mollusca) | *Chaetopleura apiculata* | Equal | 3D | Not studied | | |
| REF[8] | Errantia (Annelida) | *Alitta virens* | Unequal | 8-cell stage at 1c, 1d and 1D; all micromeres at 16-cell (no vegetal activity); 3d and 3D activity, off in micromeres; then 1q1 and 2d descendant, 3c, 3d, 3D. Fades after that. | No | 40 µM | Presumably, disrupts mesoderm bands migration to an antero-ventral position |
| REF[9] | Errantia (Annelida) | *Platynereis dumerilii* | Unequal | Late cleavage in nephroblasts, then later close to the blastopore. | No | 10, 25 or 50 µM | Shorter trunk. Reduced musculature. |
| REF[10] | Sedentaria (Annelida) | *Capitella teleta* | Unequal | The earliest detectable ERK/MAPK activation is during epiboly (gastrulation) in cells positioned around the blastopore lip. | No | 5, 10, 20 or 50 µM | Shorter and narrower posterior trunk. Reduced musculature. |
| REF[4] | Sedentaria (Annelida) | *Hydroides hexagonus* | Equal | 4d | Not studied | | |
| REF[11] | Chaetopteriformia (Annelida) | *Chaetopterus pergamentaceus* | Unequal | Not assessed | No | 20 µM | Absence of hindgut, reduced musculature. |
| This study | Palaeoannelida (Annelida) | *Owenia fusiformis* | Equal | First in four of the most animal micromeres ($1q^{111}$) at the 3q stage (4hpf), later in 4d at the coeloblastula stage (5hpf), and at 6hpf, in six additional cells $2a^1$-$2c^1$ and $2a^2$-$2c^2$ | Yes | 10 µM | Antero-ventral radialisation |

**Supplementary Table 2. di-P-ERK1/2 activation in the coeloblastula under different treatments.**

| Replicate | Treatment | di-P-ERK1/2 enrichment at the coeloblastula stage (5 hpf) | | n |
|---|---|---|---|---|
| | | yes | no | |
| 1 | BFA | 1 | 35 | 36 |
| | DMSO | 45 | 9 | 54 |
| | U0126 | 7 | 47 | 54 |
| | DMSO | 51 | 12 | 63 |
| 2 | BFA | 0 | 120 | 120 |
| | DMSO | 126 | 1 | 127 |
| | U0126 | 2 | 135 | 137 |
| | DMSO | 212 | 3 | 215 |
| 3 | BFA | 2 | 12 | 14 |
| | DMSO | 12 | 1 | 13 |
| | U0126 | 4 | 11 | 15 |
| | DMSO | 27 | 0 | 27 |
| 1 | SU5402 | 0 | 5 | 5 |
| | DMSO | 5 | 0 | 5 |
| 2 | SU5402 | 2 | 45 | 47 |
| | DMSO | 37 | 5 | 42 |

**Supplementary Table 3. Morphological markers and gene markers used for the characterisation of all phenotypes.**

| Phenotype | Morphological markers | | | | | |
|---|---|---|---|---|---|---|
| | Chaetae | Foregut | Hindgut | Muscles | Apical organ (AO) | Neurons in AO |
| **Stubby** | no | yes | yes | dorsal levators only | reduced | yes, but reduced |
| **Compressed** | yes | yes | yes | yes | reduced | yes, but reduced |
| **Radial** | no | yes | no | no | reduced | yes, but reduced |
| **Wild type** | yes | yes | yes | yes | yes | yes |
| | **Expression of gene markers in the larva** | | | | | |
| | *six3/6* | *gsc* | *cdx* | *GATA4/5/6b* | *twist* | *syt1* |
| **Stubby** | x | oral | hindgut | x | x | Apical neurons |
| **Compressed** | Oesophagus only | oral | hindgut | midgut | Trunk mesoderm | Apical neurons |
| **Radial** | Oesophagus only | Radially expanded | absent | midgut | Absent* | Apical neurons |
| **Wild type** | Apical organ and oesophagus | oral | hindgut | midgut | Trunk mesoderm | Apical neurons |

* Some windows of treatment show reduced number of *twist*+ cells

**Supplementary Table 4. Larval phenotypes at different concentrations of BFA and U0126 treatments.**

| Replicate | Treatment | Concentration (μM) | Radial n | Radial % | Wild type n | Wild type % | Total |
|---|---|---|---|---|---|---|---|
| 1 | BFA | 0.1 | 3 | 6 | 47 | 94 | 50 |
| | | 1 | 22 | 91,67 | 2 | 8,33 | 24 |
| | | 5 | 21 | 95,45 | 1 | 4,55 | 22 |
| | | 10 | 26 | 100 | 0 | 0 | 26 |
| | U0126 | 0.1 | 2 | 8 | 23 | 92 | 25 |
| | | 1 | 15 | 57,69 | 11 | 42,31 | 26 |
| | | 5 | 25 | 92,59 | 2 | 7,41 | 27 |
| | | 10 | 34 | 100 | 0 | 0 | 34 |
| 2 | BFA | 0.1 | 0 | 0 | 74 | 100 | 74 |
| | | 1 | 68 | 100 | 0 | 0 | 68 |
| | | 5 | 12 | 100 | 0 | 0 | 12 |
| | | 10 | 35 | 100 | 0 | 0 | 35 |
| | U0126 | 0.1 | 29 | 44,62 | 36 | 55,38 | 65 |
| | | 1 | 63 | 100 | 0 | 0 | 63 |
| | | 5 | 65 | 100 | 0 | 0 | 65 |
| | | 10 | 70 | 100 | 0 | 0 | 70 |
| | DMSO | 0.1 | 0 | 0 | 69 | 100 | 69 |
| | | 1 | 0 | 0 | 80 | 100 | 80 |
| | | 5 | 0 | 0 | 74 | 100 | 74 |
| | | 10 | 0 | 0 | 62 | 100 | 62 |
| | | 10 | 0 | 0 | 34 | 100 | 34 |

**Supplementary Table 5. Larval phenotypes during staggered time windows of BFA and U0126 treatments.**

| Treatment | Time window | Wild type | | Compressed | | Radial | | Total |
|---|---|---|---|---|---|---|---|---|
| | | n | % | n | % | n | % | |
| U0126 | 0.5hpf to 2c | 30 | 88,24 | 0 | 0 | 4 | 11,76 | 34 |
| | 0.5hpf to 4c | 19 | 27,14 | 0 | 0 | 51 | 72,86 | 70 |
| | 0.5hpf to 8c | 1 | 1,45 | 0 | 0 | 68 | 98,55 | 69 |
| | 0.5hpf to 3hpf | 0 | 0 | 0 | 0 | 84 | 100 | 84 |
| | 0.5hpf to 4hpf | 0 | 0 | 0 | 0 | 79 | 100 | 79 |
| | 0.5hpf to 5hpf | 0 | 0 | 0 | 0 | 1873 | 100 | 1873 |
| | 3hpf to 5hpf | 0 | 0 | 0 | 0 | 24 | 100 | 24 |
| | 4hpf to 6hpf | 0 | 0 | 0 | 0 | 517 | 100 | 517 |
| | 4hpf to 7hpf | 5 | 23,81 | 0 | 0 | 16 | 76,19 | 21 |
| BFA | 0.5hpf to 2c | 29 | 100 | 0 | 0 | 0 | 0 | 29 |
| | 0.5hpf to 4c | 60 | 100 | 0 | 0 | 0 | 0 | 60 |
| | 0.5hpf to 8c | 65 | 100 | 0 | 0 | 0 | 0 | 65 |
| | 0.5hpf to 3hpf | 39 | 43,82 | 50 | 56,18 | 0 | 0 | 89 |
| | 0.5hpf to 4hpf | 30 | 41,1 | 12 | 16,44 | 31 | 42,47 | 73 |
| | 0.5hpf to 5hpf | 0 | 0 | 0 | 0 | 795 | 100 | 795 |
| | 3hpf to 5hpf | 0 | 0 | 0 | 0 | 21 | 100 | 21 |
| | 4hpf to 6hpf | 0 | 0 | 0 | 0 | 10 | 100 | 10 |
| | 4hpf to 7hpf | 0 | - | 0 | - | 0 | - | 15 |

**Supplementary Table 6. List of candidate genes and their gene annotation IDs in differential gene expression analyses.**

| Candidate | Gene Id |
|---|---|
| *AP2* | OFUSG17466 |
| *BAMBI* | OFUSG04738 |
| *cdx* | OFUSG03177 |
| *delta* | OFUSG11344 |
| *fer3* | OFUSG04687 |
| *foxh1* | OFUSG04867 |
| *foxQ2* | OFUSG09682 |
| *gsc* | OFUSG11740 |
| *hand2* | OFUSG03211 |
| *irxA* | OFUSG05354 |
| *lhx1/5* | OFUSG14965 |
| *msx2a* | OFUSG05673 |
| *noggin* | OFUSG18439 |
| *notch-like* | OFUSG01867 |
| *POU4* | OFUSG03481 |
| *POU3* | OFUSG01141 |
| *rhox* | OFUSG21500 |
| *six3/6* | OFUSG12368 |
| *twist* | OFUSG04722 |
| *wnt1* | OFUSG16504 |
| *wnt4* | OFUSG13755 |
| *wntA* | OFUSG09779 |

**Supplementary Table 7. Candidate genes expression (ISH) in coeloblastulae and larval phenotypes in control, BFA and U0126 phenotypes treated from 0.5 hpf to 5 hpf.**

| Treatment | Stage | Gene | Wild type | | Radial | | Total | Notes |
|---|---|---|---|---|---|---|---|---|
| | | | n | % | n | % | | |
| control | Coeloblastula | *foxH* | 4 | 80 | 1 | 20 | 5 | |
| control | Larva | *foxH* | 13 | 92,86 | 1 | 7,14 | 14 | |
| U0126 | Coeloblastula | *foxH* | 0 | 0 | 9 | 100 | 9 | |
| U0126 | Larva | *foxH* | 0 | 0 | 13 | 100 | 13 | |
| control | Coeloblastula | *foxH* | 14 | 82,35 | 3 | 17,65 | 17 | |
| control | Larva | *foxH* | 21 | 100 | 0 | 0 | 21 | |
| BFA | Coeloblastula | *foxH* | 0 | 0 | 22 | 100 | 22 | |
| BFA | Larva | *foxH* | 0 | 0 | 2 | 100 | 2 | |
| control | Coeloblastula | *lhx1/5* | NA | NA | NA | NA | 59 | Faint ISH |
| control | Larva | *lhx1/5* | 43 | 100 | 0 | 0 | 43 | |
| U0126 | Coeloblastula | *lhx1/5* | NA | NA | NA | NA | 36 | Faint ISH |
| U0126 | Larva | *lhx1/5* | 0 | 0 | 116 | 100 | 116 | |

| | | | | | | | | |
|---|---|---|---|---|---|---|---|---|
| BFA | Coeloblastula | *lhx1/5* | NA | NA | NA | NA | 109 | Faint ISH |
| BFA | Larva | *lhx1/5* | 0 | 0 | 14 | 100 | 14 | |
| control | Coeloblastula | *wnt1* | NA | NA | NA | NA | 72 | Faint ISH |
| control | Larva | *wnt1* | 57 | 100 | 0 | 0 | 57 | |
| U0126 | Coeloblastula | *wnt1* | NA | NA | NA | NA | 75 | Faint ISH |
| U0126 | Larva | *wnt1* | 0 | 0 | 78 | 100 | 78 | |
| BFA | Coeloblastula | *wnt1* | NA | NA | NA | NA | 97 | Faint ISH |
| BFA | Larva | *wnt1* | 0 | 0 | 19 | 100 | 19 | |
| control | Coeloblastula | *wntA* | 73 | 100 | 0 | 0 | 73 | |
| control | Larva | *wntA* | 57 | 98,28 | 1 | 1,72 | 58 | |
| U0126 | Coeloblastula | *wntA* | 0 | 0 | 63 | 100 | 63 | |
| U0126 | Larva | *wntA* | 0 | 0 | 111 | 100 | 111 | |
| BFA | Coeloblastula | *wntA* | 0 | 0 | 71 | 100 | 71 | |
| BFA | Larva | *wntA* | 0 | 0 | 44 | 100 | 44 | |
| control | Coeloblastula | *wnt4* | NA | NA | NA | NA | 22 | Faint ISH |
| control | Larva | *wnt4* | 29 | 100 | 0 | 0 | 29 | |
| U0126 | Coeloblastula | *wnt4* | NA | NA | NA | NA | 118 | Faint ISH |
| U0126 | Larva | *wnt4* | 0 | 0 | 206 | 100 | 206 | |
| BFA | Coeloblastula | *wnt4* | NA | NA | NA | NA | 134 | Faint ISH |
| BFA | Larva | *wnt4* | 0 | 0 | 40 | 100 | 40 | |
| control | Coeloblastula | *delta* | 71 | 100 | 0 | 0 | 71 | |
| control | Larva | *delta* | 25 | 100 | 0 | 0 | 25 | |
| U0126 | Coeloblastula | *delta* | 0 | 0 | 106 | 100 | 106 | |
| U0126 | Larva | *delta* | 0 | 0 | 109 | 100 | 109 | |
| BFA | Coeloblastula | *delta* | 0 | 0 | 66 | 100 | 66 | |
| BFA | Larva | *delta* | 0 | 0 | 33 | 100 | 33 | |
| control | Larva | *six3/6* | 70 | 94,59 | 0 | 0 | 74 | |
| U0126 | Larva | *six3/6* | 0 | 0 | 40 | 100 | 40 | |
| BFA | Larva | *six3/6* | 0 | 0 | 106 | 100 | 106 | |
| control | Larva | *gata4/5/6* | 11 | 100 | 0 | 0 | 11 | |
| U0126 | Larva | *gata4/5/6* | 0 | 0 | 9 | 100 | 9 | |
| BFA | Larva | *gata4/5/6* | 0 | 0 | 13 | 100 | 13 | |
| control | Larva | *twist* | 17 | 100 | 0 | 0 | 17 | |
| U0126 | Larva | *twist* | 0 | 0 | 61 | 100 | 61 | |
| BFA | Larva | *twist* | 0 | 0 | 17 | 100 | 17 | |
| control | Larva | *cdx* | 25 | 100 | 0 | 0 | 25 | |
| U0126 | Larva | *cdx* | 0 | 0 | 27 | 100 | 27 | |
| BFA | Larva | *cdx* | 0 | 0 | 9 | 100 | 9 | |
| control | Larva | *gsc* | 23 | 100 | 0 | 0 | 23 | |
| U0126 | Larva | *gsc* | 0 | 0 | 5 | 100 | 5 | |
| BFA | Larva | *gsc* | 0 | 0 | 4 | 100 | 4 | |
| control | Coeloblastula | *notch-like* | NA | NA | NA | NA | 2 | |
| control | Larva | *notch-like* | 3 | 100 | 0 | 0 | 3 | |
| U0126 | Coeloblastula | *notch-like* | NA | NA | NA | NA | 9 | |
| U0126 | Larva | *notch-like* | 0 | 0 | 4 | 100 | 4 | |
| control | Coeloblastula | *wntA* | 4 | 100 | 0 | 0 | 4 | |
| control | Larva | *wntA* | 21 | 100 | 0 | 0 | 21 | |
| U0126 | Coeloblastula | *wntA* | 0 | 0 | 18 | 100 | 18 | |
| U0126 | Larva | *wntA* | 0 | 0 | 17 | 100 | 17 | |
| BFA | Coeloblastula | *wntA* | 0 | 0 | 46 | 100 | 46 | |

| | | | | | | | | |
|---|---|---|---|---|---|---|---|---|
| BFA | Larva | *wntA* | 0 | 0 | 5 | 100 | 5 | |
| control | Coeloblastula | *wnt4* | | 0 | 0 | 0 | 30 | |
| control | Larva | *wnt4* | | 0 | 0 | 0 | 25 | |
| U0126 | Coeloblastula | *wnt4* | 0 | 0 | 25 | 100 | 25 | |
| U0126 | Larva | *wnt4* | 0 | 0 | 16 | 100 | 16 | |
| BFA | Coeloblastula | *wnt4* | 0 | 0 | 8 | 100 | 8 | |
| BFA | Larva | *wnt4* | 0 | 0 | 4 | 100 | 4 | |
| control | Coeloblastula | *noggin* | | 0 | 0 | 0 | 11 | |
| control | Larva | *noggin* | | 0 | 0 | 0 | 41 | |
| U0126 | Coeloblastula | *noggin* | 0 | 0 | 14 | 100 | 14 | |
| U0126 | Larva | *noggin* | 0 | 0 | 3 | 100 | 3 | |
| BFA | Coeloblastula | *noggin* | 0 | 0 | 11 | 100 | 11 | |
| BFA | Larva | *noggin* | 0 | 0 | 4 | 100 | 4 | |
| control | Coeloblastula | *BAMBI* | | 0 | 0 | 0 | 13 | |
| control | Larva | *BAMBI* | | 0 | 0 | 0 | 5 | |
| U0126 | Coeloblastula | *BAMBI* | 0 | 0 | 3 | 100 | 3 | |
| U0126 | Larva | *BAMBI* | 0 | 0 | 4 | 100 | 4 | |
| BFA | Coeloblastula | *BAMBI* | 0 | 0 | 36 | 100 | 36 | |
| BFA | Larva | *BAMBI* | 0 | 0 | 5 | 100 | 5 | |
| control | Coeloblastula | *hand2* | NA | NA | NA | NA | 54 | Faint ISH |
| control | Larva | *hand2* | 51 | 100 | 0 | 0 | 51 | |
| BFA | Coeloblastula | *hand2* | NA | NA | NA | NA | 22 | Faint ISH |
| BFA | Larva | *hand2* | 0 | 0 | 13 | 100 | 13 | |
| control | Coeloblastula | *notch-like* | NA | NA | NA | NA | 52 | Faint ISH |
| control | Larva | *notch-like* | 37 | 100 | 0 | 0 | 37 | |
| U0126 | Coeloblastula | *notch-like* | NA | NA | NA | NA | 50 | Faint ISH |
| U0126 | Larva | *notch-like* | 0 | 0 | 43 | 100 | 43 | |
| control | Coeloblastula | *foxq2* | 103 | 100 | 0 | 0 | 103 | |
| control | Larva | *foxq2* | 74 | 100 | 0 | 0 | 74 | |
| U0126 | Coeloblastula | *foxq2* | 0 | 0 | 111 | 100 | 111 | |
| U0126 | Larva | *foxq2* | 0 | 0 | 87 | 100 | 87 | |
| control | Larva | *syt1* | 108 | 100 | 0 | 0 | 108 | |
| BFA | Larva | *syt1* | 0 | 0 | 9 | 100 | 9 | |
| U0126 | Larva | *syt1* | 0 | 0 | 53 | 100 | 53 | |
| control | Larva | *elav1* | 106 | 100 | 0 | 0 | 106 | |
| BFA | Larva | *elav1* | 0 | 0 | 35 | 100 | 35 | |
| U0126 | Larva | *elav1* | 0 | 0 | 88 | 100 | 88 | |
| control | Coeloblastula | *irx* | 93 | 100 | 0 | 0 | 93 | |
| control | Larva | *irx* | 62 | 100 | 0 | 0 | 62 | |
| U0126 | Coeloblastula | *irx* | 0 | 0 | 79 | 100 | 79 | |
| U0126 | Larva | *irx* | 0 | 0 | 158 | 100 | 158 | |
| BFA | Coeloblastula | *irx* | 0 | 0 | 85 | 100 | 85 | |
| BFA | Larva | *irx* | 0 | 0 | 87 | 100 | 87 | |
| control | Coeloblastula | *foxQ2* | 55 | 100 | 0 | 0 | 55 | |
| control | Larva | *foxQ2* | 14 | 100 | 0 | 0 | 14 | |
| U0126 | Coeloblastula | *foxQ2* | 0 | 0 | 40 | 100 | 40 | |
| U0126 | Larva | *foxQ2* | 0 | 0 | 38 | 100 | 38 | |
| control | Coeloblastula | *six3/6* | 52 | 100 | 0 | 0 | 52 | |
| control | Larva | *six3/6* | 15 | 100 | 0 | 0 | 15 | |
| U0126 | Coeloblastula | *six3/6* | 0 | 0 | 56 | 100 | 56 | |

| | | | | | | | | |
|---|---|---|---|---|---|---|---|---|
| U0126 | Larva | *six3/6* | 0 | 0 | 78 | 100 | 78 | |
| control | Coeloblastula | *twist* | NA | NA | NA | NA | 50 | Not expressed |
| control | Larva | *twist* | 43 | 100 | 0 | 0 | 43 | |
| U0126 | Coeloblastula | *twist* | NA | NA | NA | NA | 43 | Not expressed |
| U0126 | Larva | *twist* | 0 | 0 | 49 | 100 | 49 | |
| control | Coeloblastula | *AP2* | 20 | 100 | 0 | 0 | 20 | |
| control | Larva | *AP2* | 27 | 100 | 0 | 0 | 27 | |
| BFA | Coeloblastula | *AP2* | 0 | 0 | 112 | 100 | 112 | |
| BFA | Larva | *AP2* | 0 | 0 | 53 | 100 | 53 | |
| control | Coeloblastula | *hand2* | NA | NA | NA | NA | 48 | Faint ISH |
| control | Larva | *hand2* | 53 | 100 | 0 | 0 | 53 | |
| BFA | Coeloblastula | *hand2* | NA | NA | NA | NA | 148 | Faint ISH |
| BFA | Larva | *hand2* | 0 | 0 | 63 | 100 | 63 | |
| control | Coeloblastula | *POU3* | 45 | 100 | 0 | 0 | 45 | |
| control | Larva | *POU3* | 44 | 100 | 0 | 0 | 44 | |
| BFA | Coeloblastula | *POU3* | 0 | 0 | 63 | 100 | 63 | |
| BFA | Larva | *POU3* | 0 | 0 | 39 | 100 | 39 | |
| control | Coeloblastula | *cdx* | 69 | 100 | 0 | 0 | 69 | |
| control | Larva | *cdx* | 52 | 100 | 0 | 0 | 52 | |
| U0126 | Coeloblastula | *cdx* | 0 | 0 | 96 | 100 | 96 | |
| U0126 | Larva | *cdx* | 0 | 0 | 99 | 100 | 99 | |
| BFA | Coeloblastula | *cdx* | 0 | 0 | 64 | 100 | 64 | |
| BFA | Larva | *cdx* | 0 | 0 | 13 | 100 | 13 | |
| control | Coeloblastula | *fer3* | 56 | 100 | 0 | 0 | 56 | |
| control | Larva | *fer3* | 25 | 100 | 0 | 0 | 25 | |
| U0126 | Coeloblastula | *fer3* | 0 | 0 | 54 | 100 | 54 | |
| U0126 | Larva | *fer3* | 0 | 0 | 32 | 100 | 32 | |
| BFA | Coeloblastula | *fer3* | 0 | 0 | 94 | 100 | 94 | |
| BFA | Larva | *fer3* | 0 | 0 | 1 | 6,25 | 16 | |
| control | Coeloblastula | *foxH* | 40 | 100 | 0 | 0 | 40 | |
| control | Larva | *foxH* | 24 | 100 | 0 | 0 | 24 | |
| U0126 | Coeloblastula | *foxH* | 0 | 0 | 74 | 100 | 74 | |
| U0126 | Larva | *foxH* | 0 | 0 | 74 | 100 | 74 | |
| BFA | Coeloblastula | *foxH* | 0 | 0 | 74 | 100 | 74 | |
| BFA | Larva | *foxH* | 0 | 0 | 22 | 100 | 22 | |
| control | Coeloblastula | *lhx1/5* | NA | NA | NA | NA | 32 | Faint ISH |
| control | Larva | *lhx1/5* | 19 | 100 | 0 | 0 | 19 | |
| U0126 | Coeloblastula | *lhx1/5* | NA | NA | NA | NA | 57 | Faint ISH |
| U0126 | Larva | *lhx1/5* | 0 | 0 | 41 | 100 | 41 | |
| BFA | Coeloblastula | *lhx1/5* | NA | NA | NA | NA | 83 | Faint ISH |
| BFA | Larva | *lhx1/5* | 0 | 0 | 12 | 100 | 12 | |
| control | Coeloblastula | *rhox* | 14 | 100 | 0 | 0 | 14 | |
| control | Larva | *rhox* | 8 | 100 | 0 | 0 | 8 | |
| U0126 | Coeloblastula | *rhox* | 0 | 0 | 40 | 100 | 40 | |
| U0126 | Larva | *rhox* | 0 | 0 | 27 | 100 | 27 | |
| BFA | Coeloblastula | *rhox* | 0 | 0 | 54 | 100 | 54 | |
| BFA | Larva | *rhox* | 0 | 0 | 11 | 100 | 11 | |
| control | Coeloblastula | *wnt1* | NA | NA | NA | NA | 6 | Faint ISH |
| control | Larva | *wnt1* | 15 | 100 | 15 | 100 | 15 | |
| U0126 | Coeloblastula | *wnt1* | NA | NA | NA | NA | 75 | Faint ISH |

| | | | | | | | | |
|---|---|---|---|---|---|---|---|---|
| U0126 | Larva | *wnt1* | 0 | 0 | 46 | 100 | 46 | |
| BFA | Coeloblastula | *wnt1* | NA | NA | NA | NA | 103 | Faint ISH |
| BFA | Larva | *wnt1* | 0 | 0 | 31 | 100 | 31 | |
| control | Coeloblastula | *msx2a2* | NA | NA | NA | NA | 35 | Not expressed |
| control | Larva | *msx2a2* | 12 | 100 | 0 | 0 | 12 | |
| U0126 | Coeloblastula | *msx2a2* | NA | NA | NA | NA | 79 | Not expressed |
| U0126 | Larva | *msx2a2* | 0 | 0 | 65 | 100 | 65 | |
| BFA | Coeloblastula | *msx2a2* | NA | NA | NA | NA | 48 | Not expressed |
| BFA | Larva | *msx2a2* | 0 | 0 | 9 | 100 | 9 | |
| control | Coeloblastula | *POU4* | 49 | 100 | 0 | 0 | 49 | |
| control | Larva | *POU4* | 49 | 100 | 0 | 0 | 49 | |
| U0126 | Coeloblastula | *POU4* | 0 | 0 | 48 | 100 | 48 | |
| U0126 | Larva | *POU4* | 0 | 0 | 26 | 100 | 26 | |
| BFA | Coeloblastula | *POU4* | 0 | 0 | 44 | 100 | 44 | |
| BFA | Larva | *POU4* | 0 | 0 | 39 | 100 | 39 | |

**Supplementary Table 8. Summary of scoring counts for gene expression (ISH) in coeloblastulae and larval phenotypes in control, BFA and U0126 phenotypes treated from 0.5 hpf to 5 hpf.**

| Treatment | Stage | Total |
|---|---|---|
| control | Coeloblastula | 1252 |
| | Larva | 1330 |
| U0126 | Coeloblastula | 1378 |
| | Larva | 1873 |
| BFA | Coeloblastula | 1594 |
| | Larva | 795 |

**Supplementary Table 9. Genes expressed in the 4d blastomere or the 4d lineage in other spiralians.**

| Species | Gene/Protein | Notes | Reference |
|---|---|---|---|
| *Tritia obsoleta* | *cdx* | Other blastomere, weakly in 4d. Later in 4d lineage | REF[12] |
| | *Delta* | Also in MR and ML | REF[13] |
| | *vasa* | 4d plus other cells, and then enriched in 4d lineage | REF[14] |
| | *nanos* | 4d + 4D | REF[15] |
| *Tubifex tubifex* | *Delta* | D lineage including 4d | REF[16] |
| *Crepidula fornicata* | *vasa* | 4d | REF[17] |
| | β-catenin | 4d lineage | REF[18] |

**Supplementary Table 10. Gene expression (FISH) at 5.5 hpf in control, BFA and U0126 phenotypes treated from 0.5 hpf to 5 hpf.**

| Time window | Stage | Treatment | Gene | Wild type | | Radial | | No expression | | Total |
|---|---|---|---|---|---|---|---|---|---|---|
| | | | | n | % | n | % | n | % | |
| 0.5hpf-5hpf | CB* | control | *gsc* | 223 | 85,44 | 0 | 0 | 38 | 14,56 | 261 |
| | | U0126 | | 11 | 16,42 | 11 | 16,42 | 15 | 22,39 | 67 |
| | | BFA | | 2 | 2,13 | 0 | 0 | 92 | 97,87 | 94 |
| | | control | *AP2* | 27 | 79,41 | 0 | 0 | 7 | 20,59 | 34 |
| | | U0126 | | 84 | 10 | 0 | 0 | 78 | 92,86 | 84 |

*Coeloblastula

**Supplementary Table 11. Larval phenotypes during staggered time windows of LY411575 treatments.**

| Time window | Treatment | Wild type | | Radial | | Compressed | | Stubby | | Total |
|---|---|---|---|---|---|---|---|---|---|---|
| | | n | % | n | % | n | % | n | % | % |
| 5 hpf to 24 hpf | control | 74 | 100 | 0 | 0 | 0 | 0 | 0 | 0 | 74 |
| | LY411575 | 0 | 0 | 0 | 0 | 0 | 0 | 112 | 100 | 112 |
| 0.5 hpf to 5 hpf | control | 45 | 100 | 0 | 0 | 0 | 0 | 0 | 0 | 45 |
| | LY411575 | 68 | 100 | 0 | 0 | 0 | 0 | 0 | 0 | 68 |

**Supplementary Table 12. Gene expression (ISH) in larval phenotypes in control and LY411575 treatments.**

| Stage | Treatment | Time window | Gene | Wild type | | Stubby | | Total |
|---|---|---|---|---|---|---|---|---|
| | | | | n | % | n | % | |
| 24 hpf | control | 0.5 hpf to 5 hpf | *gsc* | 23 | 100 | 0 | 0 | 23 |
| | LY411575 | | | 37 | 100 | 0 | 0 | 37 |
| | control | 5 hpf to 24 hpf | | 18 | 100 | 0 | 0 | 18 |
| | LY411575 | | | 0 | 0 | 10 | 100 | 10 |
| | control | 0.5 hpf to 5 hpf | *cdx* | 3 | 100 | 0 | 0 | 3 |
| | LY411575 | | | 18 | 100 | 0 | 0 | 18 |
| | control | 5 hpf to 24 hpf | | 15 | 100 | 0 | 0 | 15 |
| | LY411575 | | | 0 | 0 | 27 | 100 | 27 |
| | control | 0.5 hpf to 5 hpf | *syt1* | 20 | 100 | 0 | 0 | 20 |
| | LY411575 | | | 13 | 100 | 0 | 0 | 13 |
| | control | 5 hpf to 24 hpf | | 16 | 100 | 0 | 0 | 16 |
| | LY411575 | | | 0 | 0 | 12 | 100 | 12 |

**Supplementary Table 13. Larval phenotypes during staggered time windows of SU5402 treatments.**

| Treatment | Time window | Wild type | | Radial | | Compressed | | Total |
|---|---|---|---|---|---|---|---|---|
| | | n | % | n | % | n | % | |
| control | 0.5 hpf to 7hpf | 13 | 100 | 0 | 0 | 0 | 0 | 13 |
| SU5402 | 0.5 hpf to 24hpf | 0 | 0 | 0 | 0 | 23 | 100 | 23 |
| control | | 49 | 100 | 0 | 0 | 0 | 0 | 49 |
| control | 3hpf-5hpf | 26 | 100 | 0 | 0 | 0 | 0 | 26 |
| SU5402 | | 1 | 3,2 | 0 | 0 | 30 | 96,7 | 31 |
| SU5402 | 4hpf-6hpf | 3 | 6,5 | 43 | 93,5 | 0 | 0 | 46 |
| control | | 42 | 100 | 0 | 0 | 0 | 0 | 42 |
| control | 3hpf-7hpf | 87 | 100 | 0 | 0 | 0 | 0 | 87 |
| SU5402 | | 0 | 0 | 55 | 100 | 0 | 0 | 55 |

**Supplementary Table 14. Role of FGF-MAPK and Notch-Delta signalling pathways in axial specification across bilaterians outside Spiralia.**

| Clade | Species | FGF axial | ERK axial | Notch axial | Notes | References |
|---|---|---|---|---|---|---|
| Echinodermata | *Paracentrotus lividus* | ? | Y | ? | ERK regulating nodal role in axial polarity through the ETS protein Yan/Tel | REF[19] |
| | *Lytechinus variegatus* | ? | Y | Y | Another MAPK, p38, regulating nodal role in axial polarity. Delta as a vegetal organising center. | REFs[20,21] |
| Hemichordata | *Ptychodera flava* | ? | Y | ? | ERK role in dorsal tissue formation | REF[22] |
| | *Saccoglossus kowalevskii* | N | ? | Y | Notch role in posterior elongation | REF[23] |
| Cephalochordata | | N | N | N | | |
| Tunicata | *Ciona intestinalis* | Y | Y | N | FGF-ERK role in posterior specification of epidermis | REF[24] |
| Craniata | *Xenopus tropicalis* | Y | Y | Y | FGF-ERK in dorsal mesoderm, regulation of *cdx* and communication with Delta-Notch for posterior elongation | REFs[25-27] |
| | *Danio rerio* | Y | Y | | FGF-ERK role in dorsal fate, and later in posterior elongation | REFs[28-30] |
| Arthropoda | *Parasteatoda tepidariorum* | Y | ? | Y | FGF role in dorsal-ventral through cumulus migration. Notch-Delta role in caudal specification and patterning | REFs[31,32] |
| Nematoda | *Caenorhabditis elegans* | ? | ? | Y | Notch-Delta maternal role in setting up posterior and dorsal-ventral identities | REF[33] |

**Supplementary Note 1. Multiple Protein Alignments**

==== FoxH ====

```
>Capitella_tellata_FoxH1/1-112
--------MV-AAAIWSSDRK-CLTTAEIFTQ-LRSLFVFFGS-AYTGWESSVR-
HTLSVYPCFIHSQSSRW
KVW-SVDLSLVP---ETAFVAQSKK-Q--
>Crassostrea_gigas_FoxHa/193-324
KPPYPYTGMI-IHAINSTTEK-SLTLTGIISK-LKEMFTFFKG-SYTGWRDSVR-
HNLSHNACFVKGGRSNG
NLW-HVDISKAP---INCFKLQDTP-VAR
>Danio_rerio_FoxH1/90-214
KPPYSYLAMI-AMVIQNSPEK-KLTLSEILKE-ISTLFPFFKG-NYKGWRDSVR-
HNLSSYDCFVKVLKGKG
NFW-TVEVNRIP---LELLKRQNTA-VSR
>Homo_sapiens_FoxH1/26-153
KPPYTYLAMI-ALVIQAAPSR-RLKLAQIIRQ-VQAVFPFFRE-DYEGWKDSIR-
HNLSSNRCFRKVPKAKG
NFW-AVDVSLIP---AEALRLQNTA-LCR
>Mus_muculus_FoxH1/57-184
KPPYTYLAMI-ALVIQAAPFR-RLKLAQIIRQ-VQAVFPFFRD-DYEGWKDSIR-
HNLSSNRCFHKVPKAKG
NFW-AVDVSLIP---AEALRLQNTA-LCR
>Owenia_fusiformis_FoxH1/21-146
KPPYSYLGLM-VTAIYNSPDQ-QLSLSDICRT-LENMFPFFKQ-EYRGWKDSVR-
HNLSHNTCFVKRLKKKG
NFW-AVDLDKVP---HDAFKRQDTS-VAR
>Halicryptus_spinulosus_FoxH1/113-240
KPPYSYLGLM-VMAIQASPDR-RLTLSGIHRA-LERMFSFFTS-NYSGWKDSVR-
HNLSLNNCFVKLLKAKG
NFW-TVDISKVP---LDSFKRQDTK-ESR
>Priapulus_caudatus_FoxH1/209-338
KPPFSYLGLM-VTAIQSSPSG-KLTLSGIHRA-LENMFPFFKC-EYSGWKDSVR-
HNLSLNKCFVKVLKGKG
NHW-TVNINNVP---LDAFKRQDTK-ESR
>Patinopecten/23-143 yessoensis_FoxHa
KPPYSYAGMI-IVAIMTSPNK-MLSLSEIHDY-LRNMFDFFKQ-PYLGWKDSVR-
HNLSHCKCFVKGGKGRN
NLW-TVNMAEVT---PSLFRRQQTA-IAK
>Patinopecten/36-157 yessoensis_FoxHb
KPPYTYLGLA-VMSIELSSNK-ALNLSGIVDS-LAGMFPFFRS-SYKGWRGSVR-
HTLTKYDCFIN---DGY
GEW-TVDLTKVQ---SSAFRRQETI-VAR
>Lottia_gigantea_FoxH1/19-137
KPPYSYLGMV-ALIIQCSPGR-QQSLAGIIDT-LTDMFPFFQG-EYKGWKDSVR-
HNMTNSDCFYKVEEMKR
CKW-AIDFKKLP---EDAMVRQDRR-KTT
>Branchiostoma_floridae_FoxF/46-162
KPPYSYIALI-VMAIQSSATK-RLTLSEIYQF-LQQRFPFFRG-PYQGWKNSVR-
HNLSLNECFIKLPKGKG
HYW-TIDPASEFMF-EGSFAGDAR--VRR
```

>Capitella_tellata_FoxF/91-210
KPPYSYIALI-VMAIQSAPTK-RCTLAEIYQF-LQTRFPFFRG-SYQGWKNSVR-
HNLSLNECFIKLPKGKG
HYW-TIDPAAEFMF-EGSFRRRPRG-FRR
>Lottia_gigantea_FoxF/70-188
KPPYSYIALI-VMAIQANPTR-RCTLSEIYQY-LQQKFPFFRG-TYQGWKNSVR-
HNLSLNECFIKLPKGKG
HYW-TIDPAAEFMF-EGSFRRRPRG-FRR
>Crassostrea_gigas_FoxF1/72-190
KPPYSYIALI-VMAIQATPNK-RCTLSEIYNF-LQQRFPFFRG-TYQGWKNSVR-
HNLSLNECFIKLPKGKG
HYW-TIDPAAEFMF-EGSFRRRPRG-FRR
>Novocrania_anomala_FoxF/52-169
KPPYSYIALI-VMAIQTSVTK-RCTLSEIYNF-LQQRFPFFRG-SYQGWKNSVR-
HNLSLNECFIKLPKGKG
HYW-TIDPAAEFMF-EGSFRRRPRG-FRR
>Lingula_unguis_FoxF1/b/61-180
KPPYSYIALI-VMAIQAAPTK-RCTLSEIYQF-LQQRFPFFRG-SYQGWKNSVR-
HNLSLNECFIKLPKGKG
HYW-TIDPAAEFMF-EGSFRRRPRG-FRR
>Saccoglossus_kowalevskii_FoxF/81-198
KPPYSYIALI-VMAIQSSPTK-RLTLSEIYQF-LMNRFPFFRG-PYQGWKNSVR-
HNLSLNECFIKLPKGKG
HYW-TIDPASEFMF-EGSFRRRPRG-FRR
>Danio_rerio_FoxF1/45-164
KPPYSYIALI-VMAIQSSPTK-RLTLSEIYQF-LQSRFPFFRG-SYQGWKNSVR-
HNLSLNECFIKLPKGKG
HYW-TIDPASEFMF-EGSFRRRPRG-FRR
>Homo_sapiens_FoxF1/41-159
KPPYSYIALI-VMAIQSSPTK-RLTLSEIYQF-LQSRFPFFRG-SYQGWKNSVR-
HNLSLNECFIKLPKGKG
HYW-TIDPASEFMF-EGSFRRRPRG-FRR
>Mus_muculus_FoxF1/41-159
KPPYSYIALI-VMAIQSSPSK-RLTLSEIYQF-LQARFPFFRG-AYQGWKNSVR-
HNLSLNECFIKLPKGKG
HYW-TIDPASEFMF-EGSFRRRPRG-FRR
>Homo_sapiens_FoxF2/93-212
KPPYSYIALI-VMAIQSSPSK-RLTLSEIYQF-LQARFPFFRG-AYQGWKNSVR-
HNLSLNECFIKLPKGKG
HYW-TIDPASEFMF-EGSFRRRPRG-FRR
>Mus_muculus_FoxF2/93-212
KPPYSYIALI-VMAIQSSPSK-RLTLSEIYQF-LQARFPFFRG-AYQGWKNSVR-
HNLSLNECFIKLPKGKG
HYW-TIDPASEFMF-EGSFRRRPRG-FRR
>Terebratalia_transversa_FoxF/95-214
KPPYSYIALI-VMAIQASPTK-RSTLSEIYQF-LQQRFPFFRG-TYQGWKNSVR-
HNLSLNECFIKLPKGKG
HYW-TIDPAAEFMF-EGSFRRRPRG-FRR
>Membranipora_membranacea_FoxF/79-197
KPPYSYIALI-VMAIQSTPSK-RCTLSEIYSF-LQQKFNFFRG-SYQGWKNSVR-
HNLSLNECFIKLPKGKG
HYW-TIDPSAEFMF-EGSFRRRPRG-FRR

```
>Owenia_fusiformis_FoxF1/100-222
KPPYSYIALI-VMAIQASATK-RCTLSEIYQF-LQTRFPFFRG-SYTGWKNSVR-
HNLSLNECFKKLPKGKG
HYW-TIDPAAEFMF-EGSYRRRPRG-FRR
>Dinophilus_gyrociliatus/105-229 _FoxNA
KPPYSYIALI-VMAIQASPSK-RCTLSEIYQF-LQSRFSFFRG-QYQGWKNSVR-
HNLSLNECFIKLPKGKG
HYW-TIDPTAEFMF-EGSFRRRPRG-FRR
>Drosophila_melanogaster_FoxF/304-434
KPALSYINMI-GHAIKESPTG-KLTLSEIYAY-LQKSYEFFRG-PYVGWKNSVR-
HNLSLNECFKKLPKGKG
NYW-TIDENSAHLF-EGSLRRRPRG-YRS
>Pattela_vulgata_FoxF/1-73
-----------------------------------ERFPYYHD-HKQGWQNSIR-
HNLSLNDCFVKVPRGKG
NYW-TLDPNCEEMF-NGNYRRRKRR-VKG
>Membranipora/102-225 membranacea_FoxNA
KPPYSYIALI-TMAVLQSKDK-KLTLSGICEF-IMKRFPYYRE-KFPAWQNSIR-
HNLSLNDCFVKIPR-KG
NYW-TLDPASEDMF-NGSFLRRRKR-YKR
>Prostheceraeus_vittatus_FoxNA/91-213
KPPYSYIALI-TMAVLQSPQR-KLTLSGICEF-IMTRFPYYKE-RFPAWQNSIR-
HNLSLNDCFVKVPRGKG
NYW-TLDPASEDMF-NGSFLRRRKR-YKR
>Platynereis_dum_FoxHb/111-228
KPPYPYAGII-IMAIQESPEG-MLTLAEILTS-ISEMFAHFKS-SYQGWKDSVR-
HNLSQYPCFEMVVDKKC
NKW-TVNYDHVK---TKYFACSSSS----
>Platynereis_dum_FoxHa/26-154
KPPYPYAGMA-IMAIQQSPRG-KLTLSEILAS-ISEMFTYFRC-SYQGWKDSVR-
HNLSQYACFEILPDGKC
NKW-IVNYDHVK---PKFFARNSPS----
>Convolutriloba_macropyga_FoxH1/58-182
KPPFSYQALI-IAAIWTSEAQ-QMTLREIFVF-LEKSFSFFRG-SYRGWRDSIR-
HNLTSSCVFEKVLKKKS
NFW-KVRQEHMAVV-----QKDVSA-IQQ
>Crassostrea_gigas_FoxH1/677-805
LPPYTMSSMV-IYAIQCSPQK-ALTLSEICMS-LEAMFHVYCG-NDQAWYNKVR-
NVLSKNNHFVKMRHSRK
CLW-TVDLSLVP---LSSFRKQTTR-KES
>Crassostrea_gigas_FoxHc/677-805
LPPYTMSSMV-IYAIQCSPQK-ALTLSEICMS-LEAMFHVYCG-NDQAWYNKVR-
NVLSKNNHFVKMRHSRK
CLW-TVDLSLVP---LSSFRKQTTR-KES
>Crassostrea_gigas_FoxHb/1-116
-----MSSMV-IYAIHCSPQK-TLTLSEICVS-LEAMFHVISG-NDKAWYKKVR-
NVLSKNTHFVKMRHSGK
SLW-TVDLSLVP---LTSFRKQTTR-KES

=== DELTA ===

>Pdu_delta
```

SGVFELQLLSFMNEKGLNADGNCCHGTFFKICLKHYQFAWPGTFSLIIEAWNLRMKYRVR
CETDYYGRGCTELCRSRDDRFGHYTCSGSKVCLDGWTGSYC
>Ofus_dll1
NGAFELNLQNFKNDNGFNSDGHCCNGTFFRICLTHYMFAWPGTFSLIVQAWHLFYRYRVV
CDEHYYGNKCTEYCKPRDDKNGHYTCNGEKVCLDGYTGAYC
>Hsap_dll1
SGVFELKLQEFVNKKGLLGNRNCCRGTFFRVCLKHYQFTWPGTFSLIIEALHLKYSYRFV
CDEHYYGEGCSVFCRPRDDAFGHFTCGGEKVCNPGWKGPYC
>Hsap_delta4
SGVFQLQLQEFINERGVLASGR----TFFRVCLKHFQFTWPGTFSLIIEAWHLRYSYRVI
CSDNYYGDNCSRLCKKRNDHFGHYVCQGNLSCLPGWTGEYC
>Skow_delta
SGIFQLRLASFSNDQGRNVSGQCCSVTFFRVCLKHYQFRWPGTFSLVIEAFHLHYSYRIV
CDEHYFGEECSDFCRPRDDNLGHYTCDGNKVCLPGWGGDSV
>Tcas_Delta
SGVFELRLISFDNEAGKDDKGKCCSGPRFRICLKEYQFTWPGTFSLIVEAWHLKFEYRVT
CKSHYYGKGCENLCRPRDDQFGHYSCSGERVCLAGWTGDYC
>Isca_delta
SGVFELQLRAFSNLLSQDSRGSCCSTTWFRVCLKHYQFSWPGTFSLIVEAWHAVFSFRVV
CEEHYFGADCARLCRPRDDKFGHYACNGDVVCLPGWRGDYC
>Tcas_serrate
SGFFELQVLEMANPRGELSTGECCGGTFFRLCLKEYQFRWTRSFTLILQAVDLTYRVRVK
CDSHYYNATCTKFCRPRDDKFGHYICDGDKECIEGWKGATC
>Hsap_jagged-2
MGYFELQLSALRNVNGELLSGACCDGTYVRVCLKEYQFAWPRSFTLIVEAWDLELQIRVR
CDENYYSATCNKFCRPRNDFFGHYTCDGNKACMDGWMGKEC
>Hsap_jagged-1
SGQFELEILSMQNVNGELQNGNCCGGTYFKVCLKEYQFAWPRSYTLLVEAWDFEYQIRVT
CDDYYYGFGCNKFCRPRDDFFGHYACDGNKTCMEGWMGPEC
>Pdu_jagged
SGQFQMQLVSLQNVKGELASGYCCRGTFIRFCLREYQFAWTRAFTLVFEILHMTYSLRVS
CDSNYYNTTCTKLCRPRHDAFGHYRCDGDKVCLQGWMGTNC
>Skow_jagged
SGYFELQILSVWNAAGELENGDCCDGTQVSVCLKEFQFRWMTFCTLILKILDVAYKIRIM
CDEYYYSTDCMTFCKPRDDDFGHYTCDGNKLCNTGWSGTNC
>Acali_delta
SSAPDV-------PRSRGPQWRCLFGSVFRPKVKYEEFCSQGDFSLIIEAWHLKYAFRFT
CDSNYFGAKCADLCRARDDKFGHYSCAGTKVCLDGWDGEYC
>Pdu_dlike1
DGMIKVRFKEFENKEGKAANGHCCDGHVFKICLDKPNEPLPAKVSVKVRIDDLEFEVFRE
CTHNFYGHDCSAFCQPAPRFKNQYLCDGNKICTSDWGGVDC
>Pdu_dlike3
SGEVRIKFIKYENDKGKGANGQCCDGHMFSVCLDAPGTVFPAEFDFKVVVFDLVFEASMV
CSNGYYGTNCGTHC-PAPSDTSHYECDGQKVCLQGWTGADC
>Pdu_dlike2
SGIASVKFVNYIGD-GKMADGKCCVPLFFTVCLNDPKQSWATAIDLTVTVNHLFMSISVG
CDEYYFGPGCGVYCKSSAGD--NYECGGEKICKAGWTGADC
>Smed_delta
KGVFEIHIKKYTLFPTLNLN-KCCPTIFMEICLSIYQERWKRNFNIIVKIKHLLVSFRYT
CSQNYHGEECDKLCKPSANERGFYNCSGDRICHTGYSGSYC

=== WNT ===

>Of_wnt1
QRRLVGAKVAIDECAYQFKNRRWNCGCRETAFIYAITSAAISHTVAQSCSDGSVWEWGGC
SDNADFGYDFSQDFIDVVEDLRCMMNLHNNEAGRTTVIQELRQECKCHSGSCTMKTCWKK
LAPFRVVSHKLKDRFPGVVDLVYYEKSPTFCEADNYLNSRGTRGRECNATSIGDGCDLMC
CGRGHTSETYLVRERCNCTFHWCCTVNCDICTRSRIRNTCL
>Lg_wnt1
QKKLVGAKMAIDECKYQFKNRRWNCGCRETAFIYAATSAAVSHSIARACSEGSIWEWSGC
SDNARYGHKFSRRFVDVLEDFRYMMNLHNNEAGRVHVSSGMKQECKCHSGSCTIKTCWMR
LPPFRNIGHILKDRFPGRRDLVYFEHSPTFCEKENMIGFEGTAGRECNSTSLGNGCDLMC
CGRGYKSETFPVKERCHCTFHWCCQVKCQVCTRLKVRNTCL
>Pd_wnt1
QRRVVGARVAIHECQFQFRNRRWNCGTRETAFIYAVTAAGVTHSVARACSEGSIWEWGGC
SDNIEFGQRFSREFVDLVEDLRYMMNLHNNQAGRIHVVSEQHQECKCHSGSCTVKTCWMR
LAPFRQTGARLKDRFPGPQDLVYFEESPTFCDENRTLGLQGTTGRQCNVSSIGDGCDLMC
CGRGWVEETYLSKERCNCTFHWCGQVTCHICNRTRVRHLCL
>Ct_wnt1
QRRLVAARMAVDECQRQFSTRRWNCGIRECAFIYAIMSAALAHSIARSCAEGSIWEWGGC
SDNAEFGRKFSHDFIDVAEDLKCLMNLHNSEAGRTQVSSEMSKECKCHSGSCTVKTCWMK
LPMFGRVGKVVKDRFPEPKDLVYFERSPTFCTKDPSIGHTGTHGRPCNASSIGEGCDLLC
CGRGYRSELYTARERCNCIFHWCCKVTCDTCTKTKVRHICL
>Sk_wnt1
QRKLISVGMSKTECKWQFKERRWNCGCRETAFIYAITASAVAHSVARSCSEGSIWEWGGC
SDNADFGSNFSRKFVDAGEDLRYYMNKHNNAAGRRIVTDNMRRECKCHSGSCQVKTCWMR
LPTFREVGDILKERFPTASDLVYFEESPDFCELNKKVGSLGTRGRQCNNTSIGDGCDLLC
CERGYRSEIEQVTERCSCTFHWCCQVKCETCVTQRTVHTCL
>Hs_wnt1
QRRLIGLQSAVRECKWQFRNRRWNCGCRETAFIFAITSAGVTHSVARSCSEGSIWHWGGC
SDNIDFGRLFGREFVDSGEDLRFLMNLHNNEAGRTTVFSEMRQECKCHSGSCTVRTCWMR
LPTLRAVGDVLRDRFPSPHDLVYFEKSPNFCTYSGRLGTAGTAGRACNSSSPADGCELLC
CGRGHRTRTQRVTERCNCTFHWCCHVSCRNCTHTRVLHECL
>Dm_wnt1
QRRLVGANLAISECQHQFRNRRWNCGCRETSFIYAITSAAVTHSIARACSEGTIWEWGGC
SDNIGFGFKFSREFVDTGENLREKMNLHNNEAGRAHVQAEMRQECKCHSGSCTVKTCWMR
LANFRVIGDNLKARFPGSKDLVYLEPSPSFCEKNLRQGILGTHGRQCNETSLGDGCGLMC
CGRGYRRDEVVVVERCACTFHWCCEVKCKLCRTKKVIYTCL
>Tc_wnt1
QRRLAGANNAIHECQHQFRNQRWNCGCRETAFIYAITSAAVTHAIARACSEGSIFEWGGC
SDNIGFGFTVSREFVDAGETIREKMNLHNNEAGRWHVKDQMRQECKCHSGSCTIKTCWMR
LPPFRVIGDLLKDRFPGTKDLVYYEMSPGFCEKNPKLGIQGTHGRLCNDTSMGDGCDIMC
CGRGYRTQEVVVFERCNCTFHWCCEVKCDVCRTKRTIHTCV
>Bf_wnt1
QRRLVRPMLAIKECHHQFSKWRWNCGCTQTAFIYAVMSAAVAHEVGRNCAEGTIWEWGGC
SDNVEFGKQFAKQFVDAGESVRYLVNMHNNEAGRVAVAENLRRECKCHSGSCTLKTCWMR
LPNFRDVGDSLKEKFPTDNDLVYHERSPNFCRNNPRLGFEGTRGRECNVTSRGDGCDLLC
CGRGYATRQEVTKERCNCTFQWCCQVKCEECVRTKTIHTCL
>Bf_wnt6
------------------------------------------------------WEWGGC
GDDIDFGYTKSREFMDAQTDIRTLLTLHNNEAGRLAEKNFMRTECKCHSGSCAVKTCWKK
MPIFREVGVRLKERFPTAEDLVYTNESPNFCKRNRKTGSQGTKGRACNATSMGGGCDLLC
CGRGYKERQVVVGENCKCRFHWCCVVKCSKCTAVKTVHECL
>Sk_wnt6

--------------------------------------AGVTYAVTQACSMGELWEWGGC
ADNIEFGYQKSKEFMDAQDDIRTLINLHNNEAGRLAIKKNMRIECKCHSGTCTVKTCWQK
MPVFRFVGNRLKEKFPTDEDLVYTAKSPDFCEPNRRVGSLGTGGRNCNNTSLDGGCDQLC
CGRGYKEETITVTENCKCRFHWCCVVNCDTCTTKKTIHKCN
>Lg_wnt6
QRAICGAKVALMECQYQFRTRKWNCDTREAAFVYAITAAGVVYAVTEACSMGRLWEWGGC
GDNVDFGYQKSREFMDARRDVTTLVQLHNNEAGRQAVRKYMRKECKCHSGSCTLKTCWRK
MPLFRDVGNRLKQKFPSKEDLVYSEESPDFCRRNKKEGALGTRGRECDPTSMGGGCDLLC
CGRNYSKKQVTIKENCNCRFMWCCEVICETCQKIKTETRCL
>Hs_wnt6
QAELCGARLGVRECQFQFRFRRWNCDIRETAFVFAITAAGASHAVTQACSMGELWEWGGC
GDDVDFGDEKSRLFMDARHDIRALVQLHNNEAGRLAVRSHTRTECKCHSGSCALRTCWQK
LPPFREVGARLLERFPGRADLLYAADSPDFCAPNRRTGSPGTRGRACNSSAPDSGCDLLC
CGRGHRQESVQLEENCLCRFHWCCVVQCHRCRVRKELSLCL
>Ct_wnt6
QHEMCGAKMAISECQHQFRHRRWNCDTREAAFVHSLTSAGVLYAVTQACSLGLLWEWAGC
NEDVRFGSRKAAEFLDIPPDVQGRILLHNNKAGRASVAKYQQKICKCHSGSCELKTCVLK
MPSFRDVGDRLKERFPSGENLVYLDESPSFCKPSRKQGSLGTLDRLCNPDTSTDSCDIMC
CGRGYRSYKVVVQENCRCQFKWCCKVICQTCSRTLSIHRCN
>Pd_wnt6
HRQICGVRMALDECQVQFRDRRWNCDTRETAFVYALTAAGVLYSITHACSTGQSFEWGGC
SDNVRFGYQKTRDLLDVVRDITTLVRMHNNKAGRLAVRNHVRKVCKCHSGSCTLETCWLK
MPRIHSIGRHLKQKFPDGEDLVYTTQSPDFCRVDKKQGSLGTHGRRCNPKSKADGCELMC
CGRGYIKSLRKSLQNCQCRLVWCCQVICKTCTKVETIYNCR
>Tc_wnt6
MAEICGVELGQRECQYQFRFRRWNCDTRETGFVNAVLAAGVTYQVTRACTTGELWEWEAC
GENIDFGLKKSKDFLDTRYDMKTLVKLHNYVAGRMAIKNHMRTECKCHSGSCTLKTCWRK
MPPFREVGNRLKERFPGKTGLVYSEESPHFCLPNNTLGSFGTQGRTCVETSPGEGCSILC
CGRGSRSHDETEEKNCKCKFLWCCEVKCEKCNETRTISTCL
>Dm_wnt6
LAEICGINLGFRECEFQFRNRRWNCDSRETGFVNAITAAGVTYAVTKACTMGQLWEWGGC
SDNVNFGLRHSRVFLDAKQDLGTLVKFHNNNAGRLAIRDAMRLECKCHSGSCTVKTCWLK
MPPFREVAGRLRDRYANKYQLVFADDSPDFCTPNSKTGALGTQGRECNVTSSGDRCDRLC
CNRGHTRRIVEEQTNCKCVFKWCCEVTCEKCLEHRAVNTCL
>Of_wnt2
QRDLCGAEIGLAECKDQFSTRRWNCKGREKAYIFAISSSGVMFAVTRACAKGVLFKWGGC
SHNIKYGDKFTKEWVDAQETKDGLMNTWNNEAGRKAIKASTKLICKCHSGSCSQKICWRS
MGNFKEVGSILKEKFPTKKDLVYIDDSPDFCEYNPETGSLGTKDRLCNKSSQGDGCALMC
CGRGYHTIVRTVSEECDCTFFWCCRVVCKKCQSRVEEHFCN
>Ct_wnta
--------MGIDECQHQFRDRRWNCKTREKAYIYAVSSAGVMFAVTTACAKGELFIWGGC
SHNVAFGDRFTREFVDSNENDEGLMNLWNNNAGRKAIRTSMKLLCKCHSGSCSAKICWKT
MTGFRNIGSQLKDKFPNKKDLVYLQESPDFCSSNTTIGSLGTQGRACNKTSYGDGCSLMC
CGRGYQTTLITVVEDCNCKFVWCCNVVCDECIKREERHICN
>Pd_wnta
QMELCGASTGIEECQYQFSDRRWNCETRERAYIYAVSSAGVMYSITKACAKGDLFLWGGC
SHNVKFGERFTREFVDTKEDPDGLMNIWNNGAGRKTIKSSMRLLCKCYSGSCSVKICWRT
MAPFREIGRHLKQKFPAKDELVYMEDSPDYCEYDPGIGSLGTRGRQCNRTSYGDGCSLMC
CGRGYYTTVREIKEDCNCKFHWCCRVECDKCSKKIEEHFCN
>Lg_wnta
--------MGIDECQHQFRDRRWNCKSRETGYIYAILSAGIMYSVTRACAKGDLFEWGGC
SDNIRYGSKFSKDFVDSKESEDGLMNVWNNGAGRKMVKDELELMCKCHSGSCSVKICWRK

MKSFRAIGSALKGRFPTKKDLVYLNESPDFCEHNLENGSVGTRGRECNKTSYGDGCRLMC
CGRGYYTLIKEEKDDCDCKFYWCCRVECKKCTNVKEMHYCN
>Tc_wnta
QKQLCGARLAMDECQHQFRNSRWNCRSREAAYLSAVSAASVAFAVTRACSKGDLWKWGGC
SEDIKYGEKFSRDFLDSKETADGLMNLHNNEAGRRSVKSRMVRTCKCHSGSCSMQICWRR
LPPFRKVGDALFQRYPNKTDLVYLDDSPDYCEKNETLSILGTHGRICNRTSQGDGCRLLC
CGRGYQTRVREVEEKCKCHFVWCCNVVCDICRYRREEHVCN
>Of_wnt4
QKHICGASMAIKECQFQFRNRRWNCGTREVAFVHSISSAGVSHTVTRACSSAKLFDWAGC
SDNIAYGNAFSKTFVDAREGGRSLMNLHNNEAGRQVIDDNMRVECKCHSGTCELKTCWRA
MPTFRTVGEKLKEKFHTGSDLVYLVPSPDYCEMDPHTGSIGTSGRLCSKASKADGCDLMC
CGRGFTTRVKTVTERCMCKFHWCCYVKCKECQREVEEHTCL
>Lg_wnt4
QKKLCGARVAIEECQSQFGNRRWNCGTREAAFVHAISSAGVAHSVTRACSSGTLFEWSGC
SDNIDYGMAFSKAFVDARETGRALMNLHNNEAGRKAVDANMKVECKCHSGSCEMRTCWKA
MPSFKKVGEILKEKFHTESDLVYMVASPDFCEADKKTGSLGTHGRICNKTSKADGCELLC
CGRGYRTRIVTIKERCFCKFLWCCYVKCKECTRQIEEHTCL
>Pd_wnt4
QKKICGAIEAIHECQFQFRNRRWNCGSREASFVHAISSAGVAHPVTRACSSGTMFEWAGC
SDNIAFGTAFSRTFVDARESSRVLMNLHNNEAGRKIIEENMLTQCKCHSGSCELKTCWRA
MPSFRKIGYMLKEKFHSSTDLVYLVASPDFCERDPKTGALGTHGRRCNKTSKADGCELMC
CGRGYVTKKRVIIERCHCKFHWCCYVKCQNCKREIEEHTCL
>Ct_wnt4
QKKICGAVSAIDECQYQFQNRRWNCGTREAAFVHAISSAGVTYAVTKACSSGQVFEWAGC
SDNVAYGSAFCGMFVDARESSRALMNLHNNEAGRLAVEENMKVQCKCHSGSCEMKTCWRG
LPSFREVGAMLKDRFHETQDLVYLEASPDYCISDPETGSLGTSGRTCNRSSKADGCELMC
CGRGFNVKRRVVDERCHCKFHWCCYVKCQQCKKVVDEYVCR
>Bf_wnt4
QVQICGARMSIEECQFQFRHRRWNCGTREAAFVHSISAAGVAHAVTRACSSGELFHWAGC
SDNFAFGAAFSQTFVDARESSRALMNLHNNEAGRRNLVDHMKTECKCHSGSCELKTCWRA
MPPFREVGARLKEKFHSSSDLVYLDASPDFCVRDTKVGSMGTVGRVCNKTSKADGCELLC
CGRGYNTHTREVVERCSCKFHWCCYVKCKTCRRTVEVHTCK
>Hs_wnt4
QVQMCGAQLAIEECQYQFRNRRWNCGTREAAFVYAISSAGVAFAVTRACSSGELFQWSGC
SDNIAYGVAFSQSFVDVRESSRALMNLHNNEAGRKAILTHMRVECKCHSGSCEVKTCWRA
VPPFRQVGHALKEKFHTDEDLVYLEPSPDFCEQDMRSGVLGTRGRTCNKTSKADGCELLC
CGRGFHTAQVELAERCSCKFHWCCFVKCRQCQRLVELHTCR
>Sk_wnt4
QIQVCGADMAISECQYQFKNRRWNCGTREAAFVHAISSAGVAHSVTTACSSGELFQWSGC
SDHIDYGSIFSKEFVDARENNRSLMNLHNNEAGRRTIESNMKIQCKCHSGSCETKTCWRA
LPTFREVGEKLKEKFHTDADLVYLDESPDFCENDLKSGSLGTTGRRCNKTSKADGCDLMC
CGRGYDSYTEELVERCSCKFHWCCYVKCKKCRRTVQINICR
>Bf_wnt5
QRKLCGARQGIEECQHQFRDRRWNCGSREASFTYAIAAAGVVNAVSRACREGELWLWGGC
GDDVEYGYYFAREFVDAQEHARQLMNMHNNEAGRKLTFSNARVACKCHSGSCSLKTCWQQ
LADFRTVGNLLKDKYFTDEDLVYLNKSPDYCNADPTIGSLGTHGRECNKTGLGDGCNLMC
CGRGYNTFKREKVERCNCKFHWCCYVKCKRCRSIKNVYVCK
>Hs_wnt5a
QKKLCGAKTGIKECQYQFRHRRWNCGSRETAFTYAVSAAGVVNAMSRACREGELWLWGGC
GDNIDYGYRFAKEFVDARESARILMNLHNNEAGRRTVYNLADVACKCHSGSCSLKTCWLQ
LADFRKVGDALKEKYPTTQDLVYIDPSPDYCVRNESTGSLGTQGRLCNKTSEGDGCELMC
CGRGYDQFKTVQTERCHCKFHWCCYVKCKKCTEIVDQFVCK

>Hs_wnt5b
QRKLCGAKTGIKECQHQFRQRRWNCGSRETAFTHAVSAAGVVNAISRACREGELWLWGGC
GDNVEYGYRFAKEFVDAREQGRVLMNLQNNEAGRRAVYKMADVACKCHSGSCSLKTCWLQ
LAEFRKVGDRLKEKYPTPEDLVYVDPSPDYCLRNESTGSLGTQGRLCNKTSEGDGCELMC
CGRGYNQFKSVQVERCHCKFHWCCFVRCKKCTEIVDQYICK
>Sk_wnt5
QTKLCGAKSGIEECQYQFSERRWNCASREKAFTYAISAAGVVNAISRACREGELWLWGGC
GDNVDYGYRFTKEFVDMKEHGIMKMNLHNNEAGRKAVLSMADVMCKCHSGSCSMKTCWLQ
LANFRDVGNELKERYPTAFDLVYLDPSPDYCIYDPSTGSLGTVGRLCNKTSMGDGCNLMC
CGRGYNTFQQEVVERCHCKFHWCCYVKCKRCKKIVDMHICK
>Ct_wnt5a
-----GAKIALGECQQQFHSRRWNCGSRESAFTYAIFAAGVVHAVSRSCRDGQLWLWGGC
GDNTDYGYRFAQGFVDIRELARTLMNLHNNEAGRRAVYSHTVVACKCHSGSCSLKTCWNQ
LAPFRGTGNRIKDAYPTKEDLLYLAESPDYCEADPGIGSLGTQGRQCNKHSQGDGCNLMC
CGRGYNTYKAKVSERCQCKFHWCCYVQCKTCERVVDINTCK
>Ct_wnt5b
QTKFCGAKIALGECQQQFHSRRWNCGSRESAFTYAIFAAGVVHAVSRSCRDGQLWLWGGC
GDNTDYGYRFAQGFVDIRELARTLMNLHNNEAGRRAVYSHTVVACKCHSGSCSLKTCWNQ
LAPFRGTGNRIKDAYPTKEDLLYLAESPDYCEADPGIGSLGTQGRQCNKHSQGDGCNLMC
CGRGYNTYKAKVSERCQCNFHWC------------------
>Pd_wnt5
QIKICGAQMGIRECQWQFRHRRWNCPSREAAFINAISAAGVVHTVARSCRDGELWIWGGC
GDNSEYGYRFAEGFIDARELARTLMNVHNNEAGRRAVFMHSKVACKCHSGSCSLKTCWNQ
LPSFREVGDRLKDKYPTKEDLLYLSLSPDYCLANDKTGSMGTTGRFCNKTSPGDGCTLMC
CGRGYNTYRTVVQERCQCKFHWCCYVKCKTCQREVDVHTCK
>Lg_wnt5
QIRFCGAQLGIHECQYQFRNRRWNCGNREAGFTHSISAAGVVYAISRACREGELWIWGGC
GDNIEYGYKFAKAFVDMRELSRMLMNVHNNEAGRRAVYNFAKVACKCHSGSCSLKTCWQH
LPNFREIGNRLKDRYYTKNDIIFLDDSPDYCDKNAETGSVGTAGRECDRNSQGGGCGLLC
CGRGYNTFKRKLIERCNCKFHWCCFVRCGSCERYVDVHICR
>Tc_wnt5b
QIRLCGARQALSECKHQFENRRWNCASRESAFAHALASAGVSYAVSRACRDGQLWVWGGC
GDNLEYGYKFTQNFVDVREQGRNLMNLHNNEAGRRAVIKKSKVTCKCHSGSCSLITCWQQ
LATFREIGDYLRDKYPTAYDLVYMEDSPNYCIRDERVGSLGTQGRTCNRTSQDDGCNLLC
CGRGYNTLKSTIKERCHCKFKWCCEVECKTCVRSVDVHTCK
>Ct_wnt2
QRLLCGAKKGVKQCQRQFRHHRWNCGSKEAAFVYAISSAGVVHAITRACSQGRLFDWGGC
SDNVRYGSHFARMFVDAREDARALMNLQNNRAGRRAVRRHMTLECKCHSGACTIRTCWLA
LQEFSRVGSYLKTRYTTRSDIVFFDESPDYCVQDPLAGSLGTADRECNHTSKGHGCDVMC
CGRGYDTHVVRRMRKCDCKFHWCCYVKCRECEELVKVNTCR
>Pd_wnt2
QRKLCGAKIGVKECQAQFSQYRWNCGSREAAFVYAISSSGVVNAVTRACSKGELFDWGGC
SDNVRYGAKFSRLFIDAREDARALMNLHNNRAGRRAVKKFMKLQCKCHSGSCTIRTCWLA
MQDFRRVGAFLKSKYATRSDLVYLEDSPDYCLQDTGIGSLGTAGRECNKTSLGEGCDIMC
CGRGYDVRTEQRTEKCECKFHWCCYVQCKECTKLVDVHTCK
>Lg_wnt2
QRKMCGARLGVEECQYQFQDHRWNCGTREAAFVYSISSAGVVYSITSACSKGELFDWGGC
SDSVRFGSRFSRMFVDAKEDARALMNLHNNRAGRRAVRRFRKLMCKCHSGSCTLRTCWLA
MEDFRRVGDFLKRKYPTRSDLVYFESSPDYCIKDEESGSLGTEGRECIKGSLGGGCDIMC
CGRGYDTKTIVKQEQCECKFHWCCLVKCKECSKTMDVQTCK
>Sk_wnt2
QRQLCGAKEGVKECQFQFKNNRWNCASREAAFVYAISSAGVVHAITRSCSKGELFDWGGC

SDNIKFGSDFSRHFVDAREDARALMNLHNNRAGRRAVQKNMKLECKCHSGSCTIKTCWLA
MEEFRKVGDYLRVKYPTRLDLVYFEASPDYCVADESTGSLGTAGRVCNKTSMGGGCDIMC
CGRGYDTTRAKRTTKCECKFHWCCKVICHDCTDIVDVHTCK
>Hs_wnt2
QRQLCGVAEWTAECQHQFRQHRWNCSSRESAFVYAISSAGVVFAITRACSQGEVFDWGGC
SDNIDYGIKFARAFVDAKEDARALMNLHNNRAGRKAVKRFLKQECKCHSGSCTLRTCWLA
MADFRKTGDYLWRKYPTKNDLVYFENSPDYCIRDREAGSLGTAGRVCNLTSRGDSCEVMC
CGRGYDTSHVTRMTKCGCKFHWCCAVRCQDCLEALDVHTCK
>Hs_wnt2b
QRQLCGAREWIRECQHQFRHHRWNCSSREAAFVYAISSAGVVHAITRACSQGELFDWGGC
SDNIHYGVRFAKAFVDAKEDARALMNLHNNRCGRTAVRRFLKLECKCHSGSCTLRTCWRA
LSDFRRTGDYLRRRYATRTDLVYFDNSPDYCVLDKAAGSLGTAGRVCSKTSKGDGCEIMC
CGRGYDTTRVTRVTQCECKFHWCCAVRCKECRNTVDVHTCK
>Bf_wnt3a
QIRYCGTKLGIRECQHQFRGRRWNCASREAAFVHAITSAGVAYSVTKACAEGTSWRWGGC
SEDVLFGTKFSRDFVDARIDGRSAMDRHNNEAGRQSIMKNLQLKCKCHSGSCEIKTCWWA
QPDFRTVGNVLKDKYPGKDDLIYFEVSPNFCEPNNSTGSLGTKGRECNITSQGDGCQLMC
CGRGWNTRTEMRTEKCHCQFHWCCYVTCQECQKKHQVHTCK
>Hs_wnt3
QLRFCGVKLGIQECQHQFRGRRWNCATRESAFVHAIASAGVAFAVTRSCAEGTSWKWGGC
SEDADFGVLVSREFADAREDARSAMNKHNNEAGRTTILDHMHLKCKCHSGSCEVKTCWWA
QPDFRAIGDFLKDKYPTERDLVYYENSPNFCEPNPETGSFGTRDRTCNVTSHGDGCDLLC
CGRGHNTRTEKRKEKCHCIFHWCCYVSCQECIRIYDVHTCK
>Hs_wnt3a
QLRFCGIKIGIQECQHQFRGRRWNCATRESAFVHAIASAGVAFAVTRSCAEGTAWKWGGC
SEDIEFGGMVSREFADAREDARSAMNRHNNEAGRQAIASHMHLKCKCHSGSCEVKTCWWS
QPDFRAIGDFLKDKYPTERDLVYYEASPNFCEPNPETGSFGTRDRTCNVSSHGDGCDLLC
CGRGHNARAERRREKCRCVFHWCCYVSCQECTRVYDVHTCK
>Sk_wnt3
QLRFCGAQLGIRECQNQFKGRRWNCASRETAFVHAILSAGLVHTVTRACASGELWKWGGC
SEDIRYGTRFSRDFLDPQEYARSVMNLHNNEAGRQTIAKTMETQCKCHSGSCEVKTCWKQ
QSAFKILGDLLKEKYPTSADLIYYEPSPNFCKHDPSIGSFGTEGRRCNATSNGEGCDLMC
CGRGYNTIPEEVVERCECQFIWCCKVKCKSCRRMYDLHTCK
>Bf_wnt10
------------------------------------------------------------
------------------------------------------------SGSCNLKTCWKA
TPDFREVGVILKERFPFATDLVYFDRSPDFCDRNRELETPGTRGRICNKTSTGDSCAALC
CGRGFNIFRQTRVERCNCKF---------------------
>Hs_wnt10a
QMEVCGIQIAIHECQHQFRDQRWNCGFRESAFAYAIAAAGVVHAVSNACALGKLWEWGGC
SPDMGFGERFSKDFLDSREDIHARMRLHNNRVGRQAVMENMRRKCKCHSGSCQLKTCWQV
TPEFRTVGALLRSRFASPADLVYFEKSPDFCEREPRLDSAGTVGRLCNKSSAGDGCGSMC
CGRGHNILRQTRSERCHCRFHWCCFVVCEECRITEWVSVCK
>Hs_wnt10b
QLGLCGLHIAVHECQHQLRDQRWNCGFRESAFSFSMLAAGVMHAVATACSLGKLWEWGGC
NHDMDFGEKFSRDFLDSREDIQARMRIHNNRVGRQVVTENLKRKCKCHSGSCQFKTCWRA
APEFRAVGAALRERLRLSGELVYFEKSPDFCERDPTMGSPGTRGRACNKTSRLDGCGSLC
CGRGHNVLRQTRVERCHCRFHWCCYVLCDECKVTEWVNVCK
>Lg_wnt10a
QYDLCGVQVAIHECQFQLAEYRWNCG----------------------------------
------------------------------------------------------------
------------------------------------------------------------

```
-----------------------------------------
>Pd_wnt10
QYSLCGIQVAIHECQRQFKTHRWNCGYKETAFAYAILAAGVVTQVARACSLGKLWQWKGC
DHNVEFGNAFGRKFLDSEDDFMSKVNRHNNKVGRMTVFENLRKMCKRHSGSCEMKTCWRA
APQFHVVGEVLKQKYPSKSSLVFYETSPNFCEDSSWLDSPGTRGRYCNKTSTDDNCETLC
CGRGYNTLKVTRVERCNCRFHWCCYVVCKKCLISDWVTVCK
>Ct_wnt10
QYALCGLKMAMEECQFQFREHRWNCGYRETAFASAISAAGMSAQIALACAMGNLWVWKGC
QHNVRFGDYFTRKFWGSKKNVYSEMDVHNSRAGRMIWRENVRLHCKCHSGSCEVRTCWKA
ASSFRKVGSIIKQKFIDNTDLVYFERSPNFCEPDKTLDSPGTVGRVCNSSSLHDSCDTLC
CGRGYNTVRLTKIERCQCKFRWCCDVLCKKCLMTSWVTVCK
>Lg_wnt10b
------------------------------SFAYAISSAGVTHQVSKACSMGKLFEWGGC
SHNINFGAKYASKFLDSKEDIHAQINLHNNRAGRLAIIRHVRKQCKCHSGSCELKTCWKA
APDFRAVGTILKKKYPKHLELLFYEKSPNFCDPNPLVDSPGTTGRLCNKTSGGNNCETLC
CGRGYNTLRVKRTERCHCKFFWCCYVTCKTCEYDEWVTVCK
>Sk_wnt10
QHRICGMMVAVHECSYQMRDRRWDCGYKETAFVHAIASAGVAFQVTRSCAEGRHWKWGGC
SHNVDFGEEFAQKFLDIREDVHSRINLHNNAAGRSIVSRSREKMCKCHSASCQIKTCWMS
TPKFREIGNIVKQKYAKVADLMYSERSPDFCEPSAFLGTPGTRGRSCNNTSTGDNCDSMC
CGRGYNIKSSSKVEKCNCAFHWCCFVTCDDCATSQWVNECK
>Bf_wnt7a
------------------------------------------------------------
------------------------------------------QECKCHSGSCTTKTCWTT
LPKFRELGYILKDKYPMDTDLVYIEKSPNYCEEDPVTGSVGTQGRMCNKTAQQDGCDLMC
CGRGYNTHQYSRVWQCNCKFHWCC-----------------
>Hs_wnt7a
QRAICGSQMGLDECQFQFRNGRWNCGSREAAFTYAIIAAGVAHAITAACTQGNLWKWGGC
SADIRYGIGFAKVFVDARENARTLMNLHNNEAGRKILEENMKLECKCHSGSCTTKTCWTT
LPQFRELGYVLKDKYPMDTDLVYIEKSPNYCEEDPVTGSVGTQGRACNKTAPQSGCDLMC
CGRGYNTHQYARVWQCNCKFHWCCYVKCNTCSERTEMYTCK
>Hs_wnt7b
QRAICGAQMGINECQYQFRFGRWNCGSREAAFTYAITAAGVAHAVTAACSQGNLWKWGGC
SADVRYGIDFSRRFVDARENARRLMNLHNNEAGRKVLEDRMQLECKCHSGSCTTKTCWTT
LPKFREVGHLLKEKYPMETDLVYIEKSPNYCEEDAATGSVGTQGRLCNRTSPGDGCDTMC
CGRGYNTHQYTKVWQCNCKFHWCCFVKCNTCSERTEVFTCK
>Bf_wnt7b
QRAICGAQRGIDECRYQFRHSRWNCGSKEAAFTYAISSAALVHAIVTACSQGNIWKWGGC
SADVKYGLRFCKKFVDARENARALMNLHNNEAGRKVIDQHTRLECKCHSGSCTMKTCWIT
LPRFREVGNILKEKYPREISLVYLRGSPNYCERDEATGSLGTHGRRCNRTSPYDGCDLMC
CGRGYNTHQFVKTWQCNCKFHWCCYVKCNQCSERTEEYTCK
>Sk_wnt7
QRSICGAQKGQEECRWQFRNSRWNCGNKEASFVYAINSAGVAHAITQACSQGNLWKWGGC
SADVDYGIRFSRVFVDAQENARVLTNLHNNEVGRLLLSECMDLECKCHSGSCTMKTCWTT
LPAFRTVGTTLLEKYPPRKSLVYLHKSPNYCEYDPKGGSSGTVGRRCNRTSTEDGCDLMC
CGRGYNTHQYTKTWQCNCKFHWCCFVNCIQCSERTEEYTCK
>Lg_wnt7
QRAICGAKLGLTECQYQFRFMRWNCGTKEAAFIYAMTSAGVSYAITQSCGLGSLWKWGGC
SADIKHGLRLARKFMDARENARSLMNLHNNRAGRKAVKDNMGTDCKCHSGSCTMKTCWTT
LPPFRKIGDSLKKRYPRRSHLVYLDKSPNYCDFDGKTGSLGTVGRKCNRTTKDDGCDLMC
CGRGYNTHQYTRTWQCNCKFHWCCYVNCNKCSERTEEYTCK
>Pd_wnt7
```

QRTICGASLASSECLFQFRKHRWNCGTKEAAFMYAIRAAGVAFAITQSCSSGNLFNWGGC
SVDVRYGLKFSRVFIDAREDSRSLMNLHNNRAGRKALKDLMARDCKCHSGSCTLRTCWRT
LPSFRSIGASLMRRYPRRADLVYLRHSPNYCENDPLTGSLGTRGRQCNRTSTGDGCGMLC
CGRGYNTHQFMRTWKCNCKFHWCCKVTCQNCSERSQIFTCK
>Ct_wnt7
QRTICGAKLAFEECSYQFRLHRWNCASKEAAYTSAIRSAGVSYIITQACSQGSIWKWGGC
SADIKYGLTFSRLFLDSKEDERALMNLHNNRAGRKAVKSQMDTQCKCLSGACTIKTCWTT
LPGFRSIGDHLKQKFPRKADLVYLKRSPSYCEKDEGIGSLGTTGRLCNRTANSNNCDLMC
CGRGYNTHQYTRTWQCDCKFHWCCHVTCDECTELTEEYTCK
>Tc_wnt7
QREMCGIRLATAECKYQFRHQRWNCGSREAAFTYAISSAGVAYAVTSACARGNIWKWGGC
SVDINFGMRFARKFMDAREDERSVMNLHNNKAGRKAVKMSLLTECKCHSGSCTMKTCWKT
LPGFRQVGDNLMKKYPKKSDLVFLQTSPNYCERDLAAGSLGTVGRSCNRTSRGDGCDLMC
CGRGYNTHQYTRTWQCRCKFHWCCYVDCDTCSERTEEYTCK
>Dm_wnt2
QRNMCGHQLGAQECQHQFRGHRWNCASREAAYTYAIASAGAAYAVTAACARGNIWKWGGC
SADVDFGMRYARRFMDAREDSRTLMNLHNNRAGRTLVKKMLRTDCKCHSGSCVMKTCWKS
LPPFRLVGDRLMLKYPKRMELIYLEASPNYCERSLQTGSQGTSGRTCQRTGHGQSCDLLC
CGRGHNTQHIRRTTQCRCQFRWCCEVKCDECDESYEEFTCK
>Ct_wnt16
QLRVCGARVGIEECHHQFKKERWNCGTKETAFMYAVTSAGVVHAVTKACSSGNLWKWGGC
SDNVDYGMWFAETFVDAPEDIRSLMNLQNNAVGRQVINDQMNLKCRCHSGSCAVKTCWRT
LTSFREAGNELKQKYNNGLDMVYIEDSPNYCRKNMKRGILGTKGRECERDPDADSCNTLC
CGRGYNTEVVRFVERCQCKFVWCCEVKCKICETITDKQTCK
>Lg_wnt16
QKEVCGARRGILECQKQFKFERWNCGTREAAFIYAILSAGVVHSVTQSCSAGNLFQWGGC
SDNVDYGLKFSRGFVDAPEDIRNLMNLHNNEVGRQIVEKNMQLRCRCHSGSCAVRTCFRS
LPNFRKVGLELKDKYISKQELVFVHRSPNYCKEDIKRGIFGTRGRVCNRSSPTESCDLLC
CGRGYNTQVVKYVERCHCKFFWCCYVKCKTCETMMDIHTCK
>Pd_wnt16
QIEMCGVRLGIHECQSQFRYERWNCATREAAFIHAVTAAGVVHAVTQSCSAGNLWKWGGC
SDNVDYGVWFAKTFVDAVEDIRSLVNLQNNQVGREAIAKQLHLRCRCHSGSCAVKSCWKT
MPNFNEVGKFLKKRYIKGDELVYLDPSPNYCRQNPDKGIMGTRGRECKKDSKGDSCDLLC
CGRGYNTEVIRMVERCHCKFVWCCKVKCKICETMVDKHTCK
>Sk_wnt16
QKTVCGARLGITECQKQFQRERWNCGNRETAFIYSVTTAGVVYAVTRACSAGNLWNWRGC
SDNIHYAIGFSKTFVDAPDLLRSLMNLHNNEVGRKAIEEQMDIQCRCHSGSCNVKSCWKT
MPHFPNVGDYLKTRYINTTEMVYMDQSPNYCMEDTLNGVPGTSGRECNRTSLLDSCDLLC
CGRGYNTQVVRYVERCGCKFIWCCYVKCNTCETMIDRYTCK
>Hs_wnt16
QKELCGARLGIQECGSQFRHERWNCGTKETAFIYAVMAAGLVHSVTRSCSAGNMWHWGGC
SDDVQYGMWFSRKFLDFPIKVLLAMNLHNNEAGRQAVAKLMSVDCRCHSGSCAVKTCWKT
MSSFEKIGHLLKDKYIHKDDLLYVNKSPNYCVEDKKLGIPGTQGRECNRTSEGDGCNLLC
CGRGYNTHVVRHVERCECKFIWCCYVRCRRCESMTDVHTCK
>Bf_wnt11
QTQLCAAESARKTCQEQFGNRRWNCGTKEAAYVHALSSAAVVHTVARACAAGYLYTWGGC
GDNVKFGLEFGSRFADAPMHTQTLMNLHNSEAGRLAVQNTMTTKCKCHSGSCNVKTCWKS
LADLTEISHELAEKYHGSGDLIFVENSPNYCMVNNRKGSYGTTGRLCNKTSVGDSCQTMC
CGRGYNDFTVTVTERCNCKYHWCCYVTCDQCTRTEKKYMCK
>Sk_wnt11
QKKICAAELTKQTCVKQFADRRWNCGTREAAFAYALAAAAISYSMAIECSSGAIFHWGGC
SDNVGYGMSFSARFADAPLQPESLMGLHNNQAGLLTVQENMKRKCKCHSGSCNVKTCWQS

LPEMEEIGSKLKRKYFTEQDLIYLTNSPDYCRQNEKTGSLGTVGRFCNKTSIGDGCDLMC
CGRGYKSMVITIVEQCQCRYHWCCYVKCKECSRTAEVQVCK
>Hs_wnt11
QVQLCAAREVMKACRRAFADMRWNCGTRESAFVYALSAAAISHAIARACTSGDLNRWGGC
ADNLSYGLLMGAKFSDAPMQANKLMRLHNSEVGRQALRASLEMKCKCHSGSCSIRTCWKG
LQELQDVAADLKTRYVKDSELVYLQSSPDFCMKNEKVGSHGTQDRQCNKTSNGDSCDLMC
CGRGYNPYTDRVVERCHCKYHWCCYVTCRRCERTVERYVCK
>Ct_wnt11
QIQICATRQTTDVCQYLFAQYRWNCGSREQAYVHALSAAALAQTISKACTQGATFKWGGC
GDDLRFGMIFSASFADSPFSKQAMMNLHNNNAGRKIISDSLVTDCKCHSGSCNIKTCWKA
LPDMRTVGTKIQERYFSENELIYYTKSPDYCLPDGGLGSMGTRGRECEKTNDGIGCQSMC
CGRGFTSQVVEVKHRCECKYFYCCYVECKTCTKKVEINRCR
>Pd_wnt11
QVRMCATREAVLTCQSLFADRRWNCGSREQAFVYAISSAAITHAVSRACSIGATFKWGGC
GDDLRFGLAYADLFAAPGGSKRHLVNSHNNAAGRKLISDSLQTACKCHSGSCSIKTCWKA
LPDLKELGMMLQKKYFQAEELIYYTKSPDYCLPDTTLGSLGTRGRECNKDSTSGGCRSMC
CGRGYTSHVMEIKQRCDCKYYWCCYVKCKTCTTKVEINRCR
>Lg_wnt11
QRRICASLLSVETCQNQFSDRRWNCGTREQAYVYGIASAALTHSVARACSIGVTFKWGGC
GDDLHFGMALGRAFTDASLSKKAMMNRHNFAAGRKIVESSLTTACKCHSGSCSIKTCWKS
LPDFDSIGATLKNRYIRSDELIYYTKSPDYCSPDAKSGSIGTHNRLCDKTSRGGGCDVMC
CGRGYDSFKMEVMERCECKYYWCCYVKCKTCVKTLNLSKCR
>Tc_wnt10
QLELCGIRQAVKECHYQFFNYRWNCGFRETAFAYAISSAGIAISVAKSCSRGVLWKWAGC
SHNLHYGSKFSKMFFESKEDIHSQVSLHNSKIGRMTVFANMQIKCKCHSGSCELKTCWKQ
VPNFHYVGKSLKEKFGLKNNLLFYEKSPHFCDAAPPLDVWGTSGRLCSLNATDSSCSSLC
CGRGYNLVKQRRTVSCFCVFRWCCTVECRDCIEEKWISICK
>Dm_wnt10
QVELCGLDMAIRECQIQFQWHRWNCGYRESAFAFAISAAGVAHSVARACSQGRLWKWGGC
SHNMDFGVEYSKLFLDCREDIQSKINLHNNHAGRIAVSNNMEFRCKCHSGSCQLKTCWKS
APDFHIVGKVLKHQFKLETSLFYYQRSPNFCERDLGADIQGTVGRKCNRNTTTDGCTSLC
CGRGHSQVIQRRAERCHCKFQWCCNVECEECHVEEWISICN
>Sk_wnta
QRELCGAEYGHTECQHQFRNQRWNCRSAETAFVHAVMSAGVTYSVTRACGMGILWNWDGC
NDNIGYGMYFSRDFMDAIENGLSLMNIHNNEAGRQVIKHEMYTKCKCHSGSCTSKICWRT
MRKMRDIGDILMDKYLPVTELVYLDTSPDWCEPNKKYFSHGTHGRFCNKTSRNDSCALMC
CGRGYQIMERRVEESCNCRFHWCCVVTCESCIRDEELHICN
>Dm_wnt5
QKKQCGARAAIQECQFQFKNRRWNCAAPEMAFIHALAAATVTSFIARACRDGQLWKWGGC
GDNLEFAYKFATDFIDSREKARSLMNLHNNEAGRRAVIKKARITCKCHSGSCSLITCWQQ
LSSIREIGDYLREKYPTAHDLIYLDESPDWCRNSYALHWPGTHGRVCHKNSSGESCAILC
CGRGYNTKNIIVNERCNCKFHWCCQVKCEVCTKVLEEHTCK
>Tc_wnt11
QAKLCAAVLAADTCQTVFKDRRWNCATREQAYVYAISSAALTYTMARACASGTLFQWGGC
GDNIHWGVYFAKRFIDNVEEEIAAVNLHNNRVGRRIIRESIQTQCKCHSGSCNVKTCWKG
LPPMFEIGRKLMKQYRISDHLLYLSKSPDYCTKDVKLGSFGTVGRKCNVTTNENSCRQLC
CGRGYRTLVEEKLERCHCKYYNCCYVKCKICRTKTQIYECG
>Ct_wnt9
QKKMCSVRLSVVECQANFKFDRWNCGYKETAYLHALTSSSLVHTFSRACAQGRLWLWGGC
GDNIQYGMKFARRFLRWMRDLQATADSHNSDVGIRVVRNGINKTCKCHSGSCTVQTCWRQ
LANFHEVGSDLKSKYPKKGDLVFLEKSPNYCEPTA--FGHGTTGRVCDLNK---NCDILC
CGRGYNIHTRIVDKPCHCQVIWCCHVKCQRCSVREDIYTCK

>Lg_wnt9
QKRMCATRLAVIECQYQFKHERWNCGFSETSFLYAISSAGLVHAFARACSKGVLWLWGGC
GDNLRYGLKFTRKFLKRARDVRAKVNQHNSRVGIKVVKENVNTTCKCHSGSCTVKTCWLQ
LSPFSKIGRTLKNKYPKRGNLLYMDESPSFCRRSR--YTPGTTGRTCDKDR---DCETMC
CGRGYNVKHTTVVKACKCQVFWCCHVKCKQCLKNQEIYMCK
>Hs_wnt9a
QRRMCAVSMSALECQFQFRFERWNCGFKETAFLYAISSAGLTHALAKACSAGRMWQWGGC
GDNLKYSSKFVKEFLGRRSDLRARVDFHNNLVGVKVIKAGVETTCKCHSGSCTVRTCWRQ
LAPFHEVGKHLKHKYPRTPELVHLDDSPSFCLAGR--FSPGTAGRRCHREK---NCESIC
CGRGHNTQSRVVTRPCQCQVRWCCYVECRQCTQREEVYTCK
>Hs_wnt9b
QKQLCAAHLGLLECQFQFRHERWNCGFKETAFLYAVSSAALTHTLARACSAGRMWQWGVC
GDNLKYSTKFLSNFLGSKRDLRARADAHNTHVGIKAVKSGLRTTCKCHSGSCAVRTCWKQ
LSPFRETGQVLKLRYPRSGDLVYMEDSPSFCRPSK--YSPGTAGRVCSREA---SCSSLC
CGRGYDTQSRLVAFSCHCQVQWCCYVECQQCVQEELVYTCK
>Sk_wnt9
QRRFCAVRLSTAECQYQFFNERWNCGFKETSFLYAISSAGLAHSVARSCSRGILWKWGGC
GDNLKYSQKFVQNFLQLESDLRAKAEKHNSEVGVRLLRQGFNTTCKCHSASCTTRTCWQQ
LSPFREIGNQLKEKYPRPSDLVYVDNSPHFCKKSR--FSPGTDGRRCLVGP---NCNSIC
CGRGYNIKTKVVEKSCKCRVVWCCYVECDMCSESVEIHLCK
>Pd_wnt9
QKRLCAAILSTNECKYQFRGERWNCGYRETAFLYAMSAAGLVHTLSRACSTHQLWLWGGC
DDNIQFGLRFTKRFFQAKADLKANIDRHNSRVGLKVVKNGLHKTCKCHSGSCTTQTCWHT
LSEFEEIGKILKMKYPRKRDLLFLEDSINFCDSSD--FSPGTKGRECNSKR---KCEEIC
CGRGHNLKVRMVRKPCNCTFTWCCHATCNECMVKKEMYTCK
>Bf_wnt8
PKSSFGAQTAMEECKHQFSWDRWNCANREASFVHAISAAGVMYVLTRNCSKGAFWTWGGC
SDDIAFGERISKMYSDGVEDARAAMNLHNNDVGRKAVRQTMKRVCKCHSGSCTTKTCWLQ
LADFRAIGVFLKKKYGLKKDMVFLEDSPDYCRENLTVGSRGTLGRECLRGGGKKSCKRLC
CGYVPKRITTEVTSSCNCKFHWCCSVKCSQCTKTVTKYICV
>Hs_wnt8a
PK---GAQSGIEECKFQFAWERWNCATRETSFIHAISSAGVMYIITKNCSMGDFWIWGGC
SDNVEFGERISKLFVDSLEDARALMNLHNNRAGRLAVRATMKRTCKCHSGSCSIQTCWLQ
LAEFREMGDYLKAKYSAEAELIFLEESPDYCTCNSSLGIYGTEGRECLQNSHNRSCGRLC
CGLQVEERKTEVISSCNCKFQWCCTVKCDQCRHVVSKYYCA
>Hs_wnt8b
PK---GAQSGIEECKYQFAWDRWNCANRETAFVHAISSAGVMYTLTRNCSLGDFWLWGGC
SDNVGFGEAISKQFVDALEDARAAMNLHNNEAGRKAVKGTMKRTCKCHSGSCTTQTCWLQ
LPEFREVGAHLKEKYISTRELVHLEDSPDYCLENKTLGLLGTEGRECLRRGRARSCRRLC
CGLAVEERRAETVSSCNCKFHWCCAVRCEQCRRRVTKYFCS
>Sk_wnt8
PK---GAMHGMEECRHQFLWDRWNCANRETSFVHAISSAGVMYTLTRNCSLGDFWKWGGC
SDNVNFGERVSKMFVDALVDAWAVMNLHNNEAGRKAVRQTLKRTCKCHSGSCTTQTCWKQ
LSEFRVIGDFLKRKYLSKKDLVYLEASPDYCRVNVSAGSMGTVGRECVRGTKKQSCKRLC
CGLKVKKTKVIEKSSCNCKFHWCCSVKCDECQQEVTKLTCQ
>Pd_wnt8
PK---GVERGITQCQQQFKWDRWNCVTREVAFVHAITAAGVTYTLTRNCSAGHIWLWGGC
SDNVHFGERISRLFLDSRVDARAIVHLHNNDVGRISIRRNLKLVCKCHSGSCTTKTCWQQ
LAGFKEVGIYLRRKYVKKGDLVYLAKSGNYCSLNGSAATYATLGRQCSRPKKGRSCKTLC
CGLKVSKRMTTVQRKCDCQFHWCCEVKCKLCVEDLALLTCS
>Ct_wnt8
SK---GTQLGLRECQEQFKWERWNCVTKEISFGQAITCAGVAHTITKNCSSGEFWKWGGC

```
SDNLRFGERVSKLFFDDRVDAMATVNLHNNEVGRTALKKTMELICKCHSGSCTTKTCWQH
LSDFRSVGTFLKRCYIRKTDLVYLQTSPDYCRVN---GSYATLGRQCVRSSAAKSCHRLC
CGLKVTDKVVEVTSSCKCKFKWCCEVACQQCRRKVELSTCT
>Tc_wnt8a
SKTILGAQMAMDECRNVFKWDRWNCATKEKAYTNAIITAGIIYSITRDCSQGVIWTWGGC
SDDSSFGEELVLKLLEDNEDAQAFINRHNNRIGREIIREKMLKTCKCHSGSCSFQTCWMQ
MPTFPEIAKQLRERYDVLPNLLYLEKSPDFCLSNNSTGWPGTRGRTCSRTTSAKSCRNLC
CGFRVRKQEKKVTKRCNCKFIWCCEVECDVCVEYVNEFTCH
>Dm_wnt4
QNHQCARRLATTHCEEQFRYDRWNCLYKETAFVHALTAAAMTHSIARACAEGRMFQWGGC
NDNLKHGKRVTRSFLDLRGDEVSEILRHDSEVGIEAVSSQMMDKCKCHSGSCSMKTCWKK
MADFNATATLLRQKYSQYTTLYYLETSPSYCAV--------TKDRQCLH----DNCGTLC
CGRGYTTQVVKQVEKCRCRFNRCCQLICDYCQRLENKYFCK
>Tc_wnt9
------------------RYDQWNCIYRETAFMHSMAAAALTFSIARACSEGTLWQWGGC
SDNIKFAKKFTRRFLQLRKNYQNAIVKYNSEVGIRIVIENVQVVCKCHSGSCSLKVCYKK
IQAFDYVSKQLKGLYKMGQKLVFLENSPSFCPT--------TVDRRCNNT---ENCATLC
CGRGFYTTQVHEINKCMCRWREMFNVECKYCRENKTIFRCK
>Dm_wnt8
QYTQFGLKQALDSCQQSFQWQRWNCPNREDVYVAAISMAAIVHTLTKDCANGVI------
ALNVPCAHEPTKALEQY-EGSGSGAIGHNRRVVGALLQRSLEQECRCKQGECQEEECVAV
LKPFEAIAQDLLQMYIPLDSLVFMQDSPNYCERDATGLWKGTRGRQCSKDGSGLSCQQLC
CGYRVRSQHVRTERRCNCKLVWGFRLQCDVCVQLERQYSCY

=== RHOX ===

>Nematode|ceh-45|Gsc|PRD
RRHRTIFSEEQLNILETTFTHYPDATTREELAVQCSLKEERVEVWFKNRRAKER
>Zebrafish|chr05.1|Gsc|PRD
RRHRTIFTEEQLQALEDLFNQYPDIHTREQLALKTQLREERVEVWFKNRRAKWR
>Human|GSC2|Gsc|PRD
RRHRTIFSEEQLQALEALFNQYPDVSTRERLAGRIRLREERVEVWFKNRRAKWR
>Mouse|Gsc2|Gsc|PRD
RRHRTIFSEEQLQALEALFNQYPDVGTRERLAVRIRLREERVEVWFKNRRAKWR
>Chicken|GSC2|Gsc|PRD
RRHRTIFTEEQLQALETLFNQYPDVITREHLANRIHLKEERVEVWFKNRRAKWR
>Human|GSC|Gsc|PRD
RRHRTIFTDEQLEALENLFTKYPDVGTREQLARKVHLREEKVEVWFKNRRAKWR
>Mouse|Gsc|Gsc|PRD
RRHRTIFTDEQLEALENLFTKYPDVGTREQLARKVHLREEKVEVWFKNRRAKWR
>Chicken|GSC|Gsc|PRD
RRHRTIFTDEQLEALENLFTKYPDVGTREQLARKVHLREEKVEVWFKNRRAKWR
>Zebrafish|gsc|Gsc|PRD
RRHRTIFTDEQLEALENLFTKYPDVGTREQLARKVHLREEKVEVWFKNRRAKWR
>Frog|gsc|Gsc|PRD
RRHRTIFTDEQLEALENLFTKYPDVGTREQLARRVHLREEKVEVWFKNRRAKWR
>Amphioxus|Gsc|Gsc|PRD
RRHRTIFTEEQLELLEKTFTHYPDVLLREELAMKVELKEERVEVWFKNRRAKWR
>Fruitfly|Gsc|Gsc|PRD
RRHRTIFTEEQLEQLEATFTHYPDVVLREQLALKVDLKEERVEVWFKNRRAKWR
>Honeybee|Gsc|Gsc|PRD
RRHRTIFTEEQLEQLEATFTHYPDVILREQLALQVDLKEERIEVWFKNRRAKWR
```

>Beetle|Gsc|Gsc|PRD
RRHRTIFSEEQLEQLEATFTHYPDVVLREQLALKVDLKEERVEVWFKNRRAKWR
>Human|HESX1|Hesx|PRD
RRPRTAFTQNQIEVLENVFNCYPGIDIREDLAQKLNLEEDRIQIWFQNRRAKLK
>Mouse|Hesx1|Hesx|PRD
RRPRTAFTQNQVEVLENVFNCYPGIDIREDLAQKLNLEEDRIQIWFQNRRAKMK
>Chicken|HESX1|Hesx|PRD
RRPRTAFTRNQIEVLENVFNSYPGIDIREELARKLDLEEDRIQIWFQNRRAKLK
>Frog|hesx1|Hesx|PRD
RRPRTAFTRSQIEILENVFNSYPGIDIREELAGKLALDEDRIQIWFQNRRAKLK
>Zebrafish|hesx1|Hesx|PRD
RRPRTAFSSVQIKILESVFNSYPGIDIREELAKKLQLDEDRIQIWFQNRRAKLK
>Mouse|Rhox13|Rhox|PRD
RGPPFHFAQWQVEEMESLFTQYPDLLTRGELARTLNVPEVKVKVWFTNRRAKQR
>Mouse|Rhox7|HD1|Rhox|PRD
--------------METMFTQYPDVLTREVLARSMDGSEAKVQIRFNNRRAKQR
>Human|TPRX1|Tprx|PRD
RQERTVYTESQQKVLEFYFDQYPNYDQRLNLAEMLSLREQQLQVWFKNRRAKLA
>Human|TPRXL|Tprx|PRD
RQDRTIYNWKQQEVLENHFEQYPDYDTRQELAEMLNLREYQVQVWFKNRRAKRS
>Chicken|LOC419032|Nobox|PRD
KKTRTFYSAEQLEELEKVFDRYPDNEKRREIAAVIGVTPQRIMVWFQNRRAKWR
>Zebrafish|wu:fi69e09|Nobox|PRD
KKTRTFYSTDQLEELERVFDHYPDGDKRKEIAAAIGVTPQRIMVWFQNRRAKWR
>Human|NOBOX|Nobox|PRD
KKTRTLYRSDQLEELEKIFDHYPDSDKRREIAQTVGVTPQRIMVWFQNRRAKWR
>Mouse|Nobox|Nobox|PRD
KKTRTLYRSDQLEELERIFDHYPDSDKRHEISQMVGVTPQRIMVWFQNRRAKWR
>Frog|nobox-l|Nobox|PRD
KKSRTLYSMDQLQELERLFDHYPDSEKRREIAEIIGVTPQRIMVWFQNRRAKWR
>Human|RHOXF1|Rhox|PRD
RTRRTKFTLLQVEELESVFTQYPDVPTRRELAENLGVTEDKVRVWFKNKRARCR
>Mouse|Gm14543|Rhox|PRD
RPLRDRFTEPQLQELEQVFNHYLRAEEGKQLARGMGVTEAKLQRWFKKRRVQFR
>Mouse|Rhox7|HD2|Rhox|PRD
RPLRDGFTEPQLQELEQVFNHYLRAEEGKQLARGMGVTEAKLQRWFKKRRVQFR
>Mouse|Gm7590|Rhox|PRD
RPLRDRFTEPQLQELEQVFNHCLRAEEGKQLARGMGVTEAKLQRWFKKRRVQFR
>Mouse|Gm6310|Rhox|PRD
LCLRDGFTEPQLQELEQVFNHYLRAEEGKQLARGMGVTEAKLQRWFKKRRVQFR
>Mouse|Rhox3a|Rhox|PRD
RRLHHRFTQWQLDELERIFNYFLSLEARKQLARWMGVNEAIVKRWFQKRREQYR
>Mouse|Rhox4a|Rhox|PRD
RSLHYNFQWWQLQELERIFNHFIRAEERRHLARWIGVSEARVKRWFKKRREHFR
>Mouse|Rhox8|Rhox|PRD
PRNRYRFTKFQLQELERIFNHYPSAAARRELARWIGVTESRVENWFKSRRAKYR
>Mouse|Rhox6|Rhox|PRD
RYRRTRFTHSQLHDLERLFTRYPSLRARRDLARWMGVDECDVQNWFRMRRALFQ
>Mouse|Rhox9|Rhox|PRD
RTRRTRFTHSQLRDLERLFNRFPSLRVRRDLARWMGVDESDVQEWFKMRRALFR
>Mouse|Rhox2a|Rhox|PRD
HGWQQSFNVLQLQELESIFNHYISTKEANRLARSMGVSEATVQEWFLKRREKYR

```
>Human|RHOXF2|Rhox|PRD
QPNVHAFTPLQLQELERIFEQFPSEFLRRRLARSMNVTELAVQIWFENRRAKWR
>Human|RHOXF2B|Rhox|PRD
QPNVHAFTPLQLQELECIFEQFPSEFLRRRLARSMNVTELAVQIWFENRRAKWR
>Mouse|Gm14544|Rhox|PRD
CRKQYKFTPEQLLELDRIFTQCPDTLQRKEIAKLMNVEECTVKIWFCNQRAKLR
>Mouse|Rhox11|Rhox|PRD
PRKAYRFTPGQLWELQAVFNQYPDALKRKELAGLLNVDEQKIKDWFNNKRAKYR
>Mouse|Rhox10|Rhox|PRD
RSNSKKYTNAQMCELEKAFTQYPDAHQRKALAKLIDVDECKVKAWFKYKRAKYR
>Mouse|Rhox12|Rhox|PRD
PRIQLGFTPRQLNELEDFFTKYPDALTRKNLAKHLYLAESKVQRWFKKRRAHYR
>Mouse|Rhox1|Rhox|PRD
CGLSNRFSRWQLQQLELLFTQYISAQDRKRLAVCLCVCEAKVQNWFQRRRAEYR
>New|Of_rhox
RTRRGQLSLRQKSQLEDAFCPNPTESQQRNLGKKIGITVDSVQKWFSKQHARCI
>Human|HOPX|Hopx|PRD
AETASGPTEDQVEILEYNFDKHPDSTTLCLIAAEAGLSEEETQKWFKQRLAKWR
>Mouse|Hopx|Hopx|PRD
AQTASGPTEDQVEILEYNFNKHPDPTTLCLIAAEAGLTEEQTQKWFKQRLAEWR
>Chicken|HOPX|Hopx|PRD
TEKSVTPTEEQLEILEYNFNKHPDPTTLCLIAAETGLSEEQTLKWFKQRLAEWR
>Frog|hopx|Hopx|PRD
GSDSTGLTNQQIEVLEYNFSKQPHNTSIMLIAAETGLTEEETKKWFKERLAKWR
>Amphioxus|Hopx|Hopx|PRD
PGQMSRISDDQERILEYNFTKTPDGITLEIIAAESGLSVEETAKWFQHRYALWR

=== FGF ===

>Ofus_FGFR_OFUSG19037.1/540-821
LSLGRILGEGAFGIVRQGEAVGIGGK--TTTTTVAIKSLKHDATDHEVTDLIREMEVMKV
IGRHINIINLLGCCTQ---DGPLYVIVEFAPNGNLRDFLRSRRPPNSGYEK-PM------
------------------------------------------------------------
----------ISQWQEQ---------------------------------VSIDAKPLLH
KDLLSFAYQIARGMEYLGSKLCIHRDLAARNVLVTEDYVLKIADFGLTRNIQNIDYYKKT
TDGRLPVKWMAPEALFDKKYTSKSDVWSYGVLLWEIFTLGGNPYPSVPV-EDLFKLLREG
HRMEKPPYCSLEVFNIMLDCWAQQPGYRPTFSDLVENL
>sp|P11362|FGFR1_HUMAN/478-754
LVLGKPLGEGCFGQVVLAEAIGLDKDKPNRVTKVAVKMLKSDATEKDLSDLISEMEMMKM
IGKHKNIINLLGACTQ---DGPLYVIVEYASKGNLREYLQARRPPGLEYCY-NP------
------------------------------------------------------------
--------------------------------------------------SHNPEEQLSS
KDLVSCAYQVARGMEYLASKKCIHRDLAARNVLVTEDNVMKIADFGLARDIHHIDYYKKT
TNGRLPVKWMAPEALFDRIYTHQSDVWSFGVLLWEIFTLGGSPYPGVPV-EELFKLLKEG
HRMDKPSNCTNELYMMMRDCWHAVPSQRPTFKQLVEDL
>sp|P21802|FGFR2_HUMAN/481-757
LTLGKPLGEGCFGQVVMAEAVGIDKDKPKEAVTVAVKMLKDDATEKDLSDLVSEMEMMKM
IGKHKNIINLLGACTQ---DGPLYVIVEYASKGNLREYLRARRPPGMEYSY-DI------
------------------------------------------------------------
--------------------------------------------------NRVPEEQMTF
KDLVSCTYQLARGMEYLASQKCIHRDLAARNVLVTENNVMKIADFGLARDINNIDYYKKT
TNGRLPVKWMAPEALFDRVYTHQSDVWSFGVLMWEIFTLGGSPYPGIPV-EELFKLLKEG
```

HRMDKPANCTNELYMMMRDCWHAVPSQRPTFKQLVEDL
>sp|P22607|FGFR3_HUMAN/472-748
LTLGKPLGEGCFGQVVMAEAIGIDKDRAAKPVTVAVKMLKDDATDKDLSDLVSEMEMMKM
IGKHKNIINLLGACTQ---GGPLYVLVEYAAKGNLREFLRARRPPGLDYSF-DT------
------------------------------------------------------------
--------------------------------------------------CKPPEEQLTF
KDLVSCAYQVARGMEYLASQKCIHRDLAARNVLVTEDNVMKIADFGLARDVHNLDYYKKT
TNGRLPVKWMAPEALFDRVYTHQSDVWSFGVLLWEIFTLGGSPYPGIPV-EELFKLLKEG
HRMDKPANCTHDLYMIMRECWHAAPSQRPTFKQLVEDL
>sp|P22455|FGFR4_HUMAN/467-743
LVLGKPLGEGCFGQVVRAEAFGMDPARPDQASTVAVKMLKDNASDKDLADLVSEMEVMKL
IGRHKNIINLLGVCTQ---EGPLYVIVECAAKGNLREFLRARRPPGPDLSP-DG------
------------------------------------------------------------
--------------------------------------------------PRSSEGPLSF
PVLVSCAYQVARGMQYLESRKCIHRDLAARNVLVTEDNVMKIADFGLARGVHHIDYYKKT
SNGRLPVKWMAPEALFDRVYTHQSDVWSFGILLWEIFTLGGSPYPGIPV-EELFSLLREG
HRMDRPPHCPPELYGLMRECWHAAPSQRPTFKQLVEAL
>sp|Q26614|FGFR_STRPU/639-912
LTVGKTIGEGAFGKVVIGEAVGIVCQ--EKTSTVAVKMLKANAMDREFSDLISELAMMKM
IGKNPNIINLLGCCTQ---EGPPYVIVEFAHHGNLRDFLRSRRPPE-EYEK-SI------
------------------------------------------------------------
--------------------------------------------------LLTTSQTLTN
KDLMSMAYQVARGMDFLASKKCIHRDLAARNVLVTEDFEMKICDFGLARDIHYIDFYRKT
TDGRLPVKWMAPEALFDRMFTTQSDVWSFGILLWEIMTLGGTPYPSVPV-EQMFDYLRSG
KRLEKPQNTSLEIYHILCECWRTSPGQRPTFCELVEDL
>sp|Q86PM4|FGFR_HYDVU/474-743
LETDCLLGEGAFGRVFRATARDLPNH--TGVQTVAVKMLKEDCCEQDLKDFISEIEVMKS
IGKHINILNLLAVSSQ---QGKLYIVVEYCRHGNLRSFLKDNRPVMQA------------
------------------------------------------------------------
--------------------------------------------------NSVITKKITL
YDLTSFCLQVARGMNFLASKKCIHRDIAARNVLVGEGYLMKIADFGLARDIHEQDYYRKC
TDGRLPVKWMAIEALFDRVYTTQSDIWSFGILAWEIVTFGGSPYPGIAL-EKLFDLLKQG
YRMERPLNCTDDMYTLMLNCWKEIPSKRPTFSQLIEDL
>sp|Q07407|FGFR1_DROME/416-688
LVLGATLGEGAFGRVVMAEV---------NNAIVAVKMVKEGHTDDDIASLVREMEVMKI
IGRHINIINLLGCCSQ---NGPLYVIVEYAPHGNLKDFLYKNRPFGRDQDR-DSS-----
------------------------------------------------------------
--------------QPP-----------------------------------PSPPAHVITE
KDLIKFAHQIARGMDYLASRRCIHRDLAARNVLVSDDYVLKIADFGLARDIQSTDYYRKN
TNGRLPIKWMAPESLQEKFYDSKSDVWSYGILLWEIMTYGQQPYPTIMSAEELYTYLMSG
QRMEKPAKCSMNIYILMRQCWHFNADDRPPFTEIVEYM
>sp|Q09147|FGFR2_DROME/712-996
LSLGSILGEGAFGRVVMAEAEGLPRSPQLAETIVAVKMVKEEHTDTDMASLVREMEVMKM
IGKHINIINLLGCCSQ---GGPLWVIVEYAPHGNLKDFLKQNRPGAPQRRS-DSD-----
------------------------------------------------------------
-----------GYLDDK-----------------------------------PLISTQHLGE
KELTKFAFQIARGMEYLASRRCIHRDLAARNVLVSDGYVMKIADFGLARDIQDTEYYRKN
TNGRLPIKWMAPESLQEKKYDSQSDVWSYGVLLWEIMTYGDQPYPHILSAEELYSYLITG
QRMEKPAKCSLNIYVVMRQCWHFESCARPTFAELVESF
>sp|Q4H3K6|FGFR_CIOIN/377-656
ILLHERIDEGFFGQVFRADLIRCAGGR-KEKVDAAVKMLKSTRTEKDMLDLLTEMDQMKR
VGKHKNIVNLLGVCTQ---NGILWLVTEYAQKGNLRDYLRRNRPSELQYEL-ST-----P

```
------------------------------------------------------------
--------------DSP---------------------------------APPRDEPLTL
RALMSASHQVARGMEYLSQKKCIHRDLAARNVLVANDFVMKIADFGLARDIRSNDYYRKE
TRGHLPYKWMALEAMTDNMFTHATDVWSFGILLWEIFSLGGSPYPGVKT-HDLVRFLRNG
DRLEQPQFASSELYRLMRDCWEESPRRRPQFRQLVEDL
>sp|P26619|PGFRA_XENLA/595-950
LVLGRILGSGAFGKVVEGAAYGLSRS--QPVMKVAVKMLKPTARSSEKQALMSELKIMTH
LGAHLNIVNLLGACTK---SGPIYIITEYCFYGDLVNYLHKNRDNFQSRHP-EK-----P
KKDLDIFGLN-------------PADESTRSYVILSFENNGDYMDMKQADTMQYVPMLEM
KEPSKYSDIQRSLYDRPASYKKKPL--SEVKNIL----------------SDDGFEGLTV
LDLLSFTYQVARGMEFLASKNCVHRDLAARNVLLAHGKIVKICDFGLARDIMHDSNYVSK
GSTFLPVKWMAPESIFDNLYTTLSDVWSFGILLWEIFSLGGTPYPGMIVDSTFYNKIKSG
YRMAKPDHATHEVYDIMVKCWNSEPEKRPSFRHLSDIV
>sp|Q9PUF6|PGFRA_CHICK/593-950
LVLGRILGSGAFGKVVEGTAYGLSRS--QPVMKVAVKMLKPTARSSEKQALMSELKIMTH
LGPHLNIVNLLGACTK---SGPIYIITEYCFYGDLVNYLHKNRDNFLSRHP-EK-----P
KKDLDIFGMN-------------PADESTRSYVILSFENTGEYMDMKQADTTQYVPMLER
KEGSKYSDIQRSVYDRPASYKKKSLSESEVKNLL----------------SDDGSEGLSL
LDLLSFTYQVARGMEFLASKNCVHRDLAARNVLLAQGKIVKICDFGLARDIMHDSNYVSK
GSTFLPVKWMAPESIFDNLYTTLSDVWSYGILLWEIFSLGGILYPGMMVDSTFYNKIKSG
YRMAKPDHATNEVYEIMVKCWNNEPEKRPSFYHLSEIV
>sp|P16234|PGFRA_HUMAN/593-950
LVLGRVLGSGAFGKVVEGTAYGLSRS--QPVMKVAVKMLKPTARSSEKQALMSELKIMTH
LGPHLNIVNLLGACTK---SGPIYIITEYCFYGDLVNYLHKNRDSFLSHHP-EK-----P
KKELDIFGLN-------------PADESTRSYVILSFENNGDYMDMKQADTTQYVPMLER
KEVSKYSDIQRSLYDRPASYKKKSMLDSEVKNLL----------------SDDNSEGLTL
LDLLSFTYQVARGMEFLASKNCVHRDLAARNVLLAQGKIVKICDFGLARDIMHDSNYVSK
GSTFLPVKWMAPESIFDNLYTTLSDVWSYGILLWEIFSLGGTPYPGMMVDSTFYNKIKSG
YRMAKPDHATSEVYEIMVKCWNSEPEKRPSFYHLSEIV
>sp|P09619|PGFRB_HUMAN/600-958
LVLGRTLGSGAFGQVVEATAHGLSHS--QATMKVAVKMLKSTARSSEKQALMSELKIMSH
LGPHLNVVNLLGACTK---GGPIYIITEYCRYGDLVDYLHRNKHTFLQHHS-DKRRP--P
SAELYSNAL--------------PVGLPLPSHVSLTGESDGGYMDMSKDESVDYVPMLDM
KGDVKYADIESSNYMAPYDNYVPSAPERTCRAT-----------------LINESPVLSY
MDLVGFSYQVANGMEFLASKNCVHRDLAARNVLICEGKLVKICDFGLARDIMRDSNYISK
GSTFLPLKWMAPESIFNSLYTTLSDVWSFGILLWEIFTLGGTPYPELPMNEQFYNAIKRG
YRMAQPAHASDEIYEIMQKCWEEKFEIRPPFSQLVLLL
>sp|P17948|VGFR1_HUMAN/827-1154
LKLGKSLGRGAFGKVVQASAFGIKKS--PTCRTVAVKMLKEGATASEYKALMTELKILTH
IGHHLNVVNLLGACTK--QGGPLMVIVEYCKYGNLSNYLKSKRDLFFLNKD-AA-----L
HMEPKKEKME----------------------------PGLEQGKKPRLDSVTSSESFAS
SGFQED----------------KSLSDVEEEEDS----------------DGFYKEPITM
EDLISYSFQVARGMEFLSSRKCIHRDLAARNILLSENNVVKICDFGLARDIYKNPDYVRK
GDTRLPLKWMAPESIFDKIYSTKSDVWSYGVLLWEIFSLGGSPYPGVQMDEDFCSRLREG
MRMRAPEYSTPEIYQIMLDCWHRDPKERPRFAELVEKL
>sp|Q8AXB3|VGFR4_DANRE/809-1135
LRLGKTLGHGAFGKVVEASAFGIDKI--STCKTVAVKMLKVGATNNEWRALMSELKILIH
IGHHLNVVNLLGACTK--RGGPLMIIVEFCKYGNLSNYLRSKRGDFVVYKSQDGKA---V
RSSSGC---------------------------------DLSELIKRRLESVASTGSSAS
SGFIED----------------KSYCDSEEEEEEQ---------------EDLYKKVLTL
EDLICYSFQVAKGMEFLASRKCIHRDLAARNILLSENNVVKICDFGLARDVYKDPDYVRK
GDARLPLKWMAPEAIFDKIYTTQSDVWSFGVLMWEIFSLGASPYPGLHIDEEFCCRLKEG
```

TRMKAPEYSSSEIYQTMLDCWHGEPSQRPTFTELVERL
>sp|P35968|VGFR2_HUMAN/834-1160
LKLGKPLGRGAFGQVIEADAFGIDKT--ATCRTVAVKMLKEGATHSEHRALMSELKILIH
IGHHLNVVNLLGACTK--PGGPLMVIVEFCKFGNLSTYLRSKRNEFVPYKT-KG-----A
RFRQGKDYVG-----------------------------AIPVDLKRRLDSITSSQSSAS
SGFVEE----------------KSLSDVEEEEAP----------------EDLYKDFLTL
EHLICYSFQVAKGMEFLASRKCIHRDLAARNILLSEKNVVKICDFGLARDIYKDPDYVRK
GDARLPLKWMAPETIFDRVYTIQSDVWSFGVLLWEIFSLGASPYPGVKIDEEFCRRLKEG
TRMRAPDYTTPEMYQTMLDCWHGEPSQRPTFSELVEHL
>sp|Q5GIT4|VGFR2_DANRE/843-1171
LKLGEPLGRGAFGQVVEATAYGIEKA--TTCTTVAVKMLKEGATSSEYRALMSELKILIH
IGHHLNVVNLLGACTK--QGGPLMVIVEYCKHGNLSSYLKSKRGEYSPYKK-RT-----P
RMPNRREVQQDE----------DP------------REGDLGLGTSTRLDICTGTAVCTR
TGEQTY----------------KTLQDEQESSDW---------------------DHLTM
EDLISYSFQVAKGMEFLASRKCIHRDLAARNILLSENSVVKICDFGLARDVYKDPDYVRK
GDARLPLKWMAPETIFDRVYTTQSDVWSFGVLLWEIFSLGASPYPGVCIDESFCRRLKEG
TRMRAPDYATPEIYQTMLDCWLDRPLDRPTFTQLVEHL
>sp|P35916|VGFR3_HUMAN/845-1169
LHLGRVLGYGAFGKVVEASAFGIHKG--SSCDTVAVKMLKEGATASEHRALMSELKILIH
IGNHLNVVNLLGACTK--PQGPLMVIVEFCKYGNLSNFLRAKRDAFSPCAE-KS-----P
EQRGFRAMV------------------------------EL--ARLDRRRPGSSDRVLFA
RFSKTE----------------GGARRASPDQEA----------------EDLWLSPLTM
EDLVCYSFQVARGMEFLASRKCIHRDLAARNILLSESDVVKICDFGLARDIYKDPDYVRK
GSARLPLKWMAPESIFDKVYTTQSDVWSFGVLLWEIFSLGASPYPGVQINEEFCQRLRDG
TRMRAPELATPAIRRIMLNCWSGDPKARPAFSELVEIL
>sp|Q5MD89|VGFR3_DANRE/866-1177
LRLGKVLGHGAFGKVIEASIFGHDKK--SSANTVAVKMLKEGATASEHKALMSELKILIH
IGNHLNVVNLLGACTK--PNGPLMVIVEYCKYGNLSNFLRAKREFFLPYRD-RS-----P
KTQSQVRRMI-----------------------------EAGQASQSEHQPSTSS-----
--------------------------TNPPRVTV----------------DDLWKTPLTI
EDLICYSFQVARGMEFLASRKCIHRDLAARNILLSENNVVKICDFGLARDIYKDPDYVRK
GNARLPLKWMAPESIFDKVYTSQSDVWSFGVLLWEIFSLGASPYPGIQIDEDFCKRLKDG
TRMRAPDNASPEIYGIMLACWQGEPRERPTFPALVEIL
>Ofus_VEGFROFUSG03166.1/880-1201
LNFGMVLGAGAFGRVVKADAVGLNDY--ESSTPVAVKMVKDCTDYDQMKSLMSELKIMIH
IGSHLNVLNLLGAVTKDIASGELYIMMEYCPHGNLRNYLLKHRATFRDVTD-DEDVFEMP
KK-----------------------------------PGADQNLSPAQLKKLEAEG-GAT
GGPSDD----------------LHYVNVP---------------------ESHREPPVTT
KDLICFSYQIARGLEYLASRKFVHRDIAARNILVAEDRVMKIADFGLAKDVYKYEEYVKK
GAGALPIKWLALESLTHKVFSVKTDVWAYGILLYETFSLGGTPYPGVDLDATFIEKLKNG
YRMEKPQFASFQIYQLMKMCWEAEPEDRPTFSEITERV
>tr|Q95P10|Q95P10_DROME/867-1261 PDGF/VEGF
LKLGKQLGAGAFGVVLKGEAKGIRRE--EPTTTVAVKMVKATADNEVVRALVSELKIMVH
LGQHLNVVNLLGAVTKNIAKRELMVIVEYCRFGNIQNFLLRNRKCFINQINPDTDHID-P
SIMTQRMSDNYELHRDTNGGGLKYANVGFPIHSYINEPHNNNTQPPTHRRNSDNDPRSGT
RAGRTGSGTATYSYDRQMDTCATVMTTVPEDDQIMSNNSVQPAWRSNYKTDSTEAMTVTT
VDLISWAFQVARGMDYLSSKKVLHGDLAARNILLCEDNVVKICDFGLARSMYRGDNYKKS
ENGKLPIKWLALESLSDHVFSTYSDVWSYGIVLWEMFSLAKVPYPGIDPNQELFNKLNDG
YRMEKPKFANQELYEIMLECWRKNPESRPLFAELEKRF
